# Omics-scale decoding of protein *N*-glycosylation using protein language model embeddings

**DOI:** 10.64898/2026.11.02.763593

**Authors:** Xiao Zhou, Xingyue Guan, Yao Zhou, Jie Wang, Zhiyu Peng, Weitong Ren, Wei Wang, Ran Xie, Wenfei Li

## Abstract

Protein language models (pLMs) learn rich sequence semantics from vast volume of protein sequences, yet the information encoded in their high-dimensional embeddings remains largely a black box, posing an interpretability challenge. Here we show that semantic features learned by pLMs can be decoded to reveal biologically meaningful signals using coevolutionary analysis without downstream supervised-training. Building on this insight, we developed MaskNGly, a zero-shot framework that identifies protein *N*-glycosylation sites directly from PLM-embeddings while achieving performance comparable to state-of-the-art supervised predictors. MaskNGly is computationally efficient and intrinsically transferable, enabling omics-scale characterization of protein glycosylation landscape. Applied across proteomes, it reveals that canonical N-X-S/T motifs gain higher glycosylation propensities with increasing organismal complexity and that glycosylation preferentially occurs at structured, solvent-exposed regions. Mutations altering high-scoring motifs are more frequently disease-associated. The framework also captures signatures of glycosylation-mediated protein allostery. These results establish an interpretable and scalable route to study glycobiology from pLM-embeddings.

## Introduction

Protein *N*-linked glycosylation (*N*-glycosylation) is one of the most prevalent and biologically important post-translational modifications (PTMs), playing essential roles in protein homeostasis [1, 2], immune regulation, and signal transduction [3, 4]. Aberrant glycosylation is closely associated with the pathogenesis of numerous cancers, neurodegenerative disorders, and autoimmune diseases [5–7]. Most *N*-glycosylation dominantly occurs at canonical N-X-S/T motifs (where X ̸= P) [8, 9]. However, only part of these motifs in natural proteins are actually glycosylated, as glycosylation modification depends strongly on local geometric and physicochemical features encoded by the full protein sequence [8, 10]. These multilayered determinants make accurate prediction of *N*-glycosylation sites a challenging task. Developing computational frameworks capable of predicting glycosylation sites and elucidating the principles governing *N*-glycosylation is therefore essential for understanding the molecular mechanisms underlying cellular function and disease, and for guiding therapeutic design.

Given the biological importance of protein glycosylation, developing algorithms capable of accurately predicting glycosylation sites has long been a major research objective, and substantial progresses have been made [11, 12]. Among these efforts, machine learning approaches achieved notable improvements of prediction accuracy by incorporating carefully selected protein features as inputs. More recently, advances in protein language models (pLMs), such as ESM-2 [13], ProtT5 [14], and ProteinBERT [15], have enabled high dimensional sequence embeddings that capture complex semantic patterns within protein sequences. Leveraging these embeddings as sequence features, machine-learning or deep neural network models have achieved further gains in prediction accuracy [16–20]. Despite these advances, supervised learning approaches face major challenges in glycosylation site prediction. They depend on precise annotation of both positive and negative sites during model training. However, experimental detection of glycosylation is often incomplete due to the low interpretation rates of glycopeptide spectra from mass spectrometry [21], making reliable negative datasets difficult to construct. The absence of detectable glycosylation signals in experiments does not necessarily mean that a given site is not a glycosylation site. Likewise, existing annotations in glycosylation datasets are subject to continual updates and corrections as quality-control stringency increases. Such label uncertainty can undermine confidence in predictions from supervised models. These limitations underscore the need for a new computational framework that is interpretable, does not rely on annotated training datasets, and is broadly applicable across species and omics-scale analysis.

On the other hand, pLMs trained on vast collections of evolutionarily diverse protein sequences learn rich and multifaceted sequence semantics. Although embeddings generated by these models have been widely applied to structure-and function-prediction tasks, they are typically treated as abstract feature representations for downstream deep-learning models, leaving their biological meaning largely opaque. An urgent need in computational biology is therefore to open this black box and directly extract biologically interpretable information from pLM embeddings. Emerging evidence suggests that pLMs can implicitly learn signals related to PTM [22]. Given the major challenges faced by supervised approaches in characterizing glycosylation landscape, three fundamental questions arise: (1) Do pLM embeddings already encode latent information relevant to *N*-glycosylation? (2) If so, can glycosylation sites be decoded directly from pLM embeddings without relying on annotated training data, and (3) how does *N*-glycosylation vary across proteomes?

In this study, we show that sequence information specifying the *N*-glycosylation sites of a protein is fully encoded within the pLM embedding (Figure 1). By directly decoding this information through a mask-and-probe based coevolutionary analysis without relying on annotated training datasets, we achieve a performance comparable to the state-of-the-art supervised learning approaches in predicting protein *N*-glycosylation sites. This annotation-free framework is intrinsically transferable, which enables omics-scale characterization of *N*-glycosylation landscape and uncovers key features of glycosylation across species, structural contexts, and disease-associated mutations. We find that the *N*-glycosylation probabilities of canonical N-X-S/T motifs increase with organismal complexity, and that glycosylation preferentially occurs at structured and solvent-exposed regions. Mutations introducing or disrupting high-scoring motifs are more frequently disease-associated. We also demonstrated that the information of glycosylation induced allosteric coupling can be inferred from the coevolutionary analysis of ESM-2 embedding, which otherwise often relies on long-timescale molecular dynamics (MD) simulations. These findings establish a training-free and interpretable framework for glycosylation prediction and open avenues for interpreting glycan-mediated biological mechanisms through protein language models.

**Figure 1.**
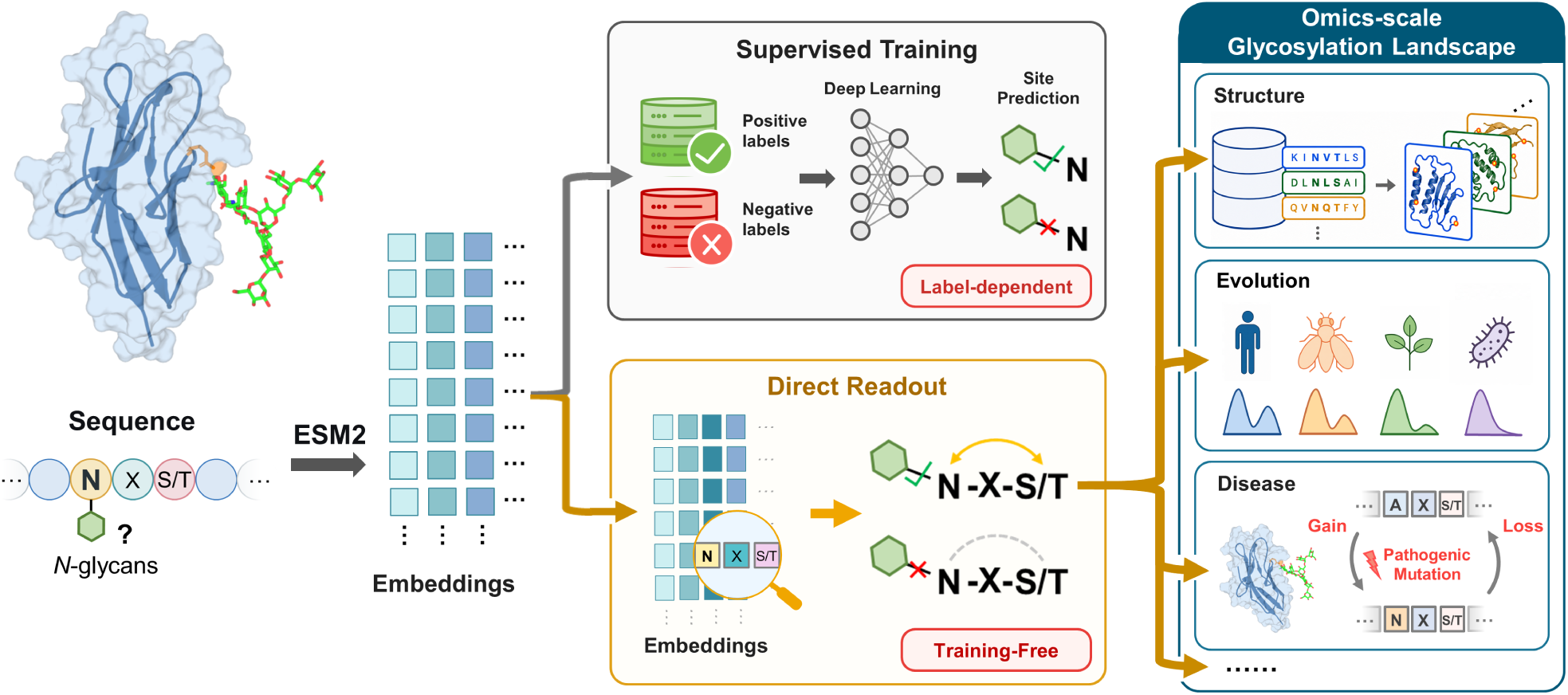
Schematic illustration of two complementary routes for predicting *N*-glycosylation sites from pLM embeddings: a supervised deep-learning approach trained on curated glycosylation annotations, and a training-free direct readout approach based on coevolutionary signals extracted from pLM embeddings.

## Results

### Quantifying coevolution by pLM embedding

Evolutionary patterns encoded within protein family sequences reflect the energy landscape underlying protein folding and function [23–26]. Leveraging these signals, deep learning methods have transformed the prediction of protein three-dimensional structures and contact maps [27–30]. In parallel, protein language models (pLMs), trained on hundreds of millions of natural protein sequences using masked language modeling [13, 31], learn representations that capture long-range dependencies and evolutionary constraints from sequence context. Previous studies have shown that protein language model ESM-2 implicitly encodes pairwise residue coevolutionary couplings within high-dimensional sequence representations [13, 32]. Extracting such couplings often requires supervised decoding, for example, by training a “Contact Head” neural network model in the structure-prediction task [13, 31]. Interestingly, recent work by Zhang et al. demonstrated that coevolutionary signals can be inferred directly through a “categorical Jacobian” calculation from ESM-2 logits, enabling unsupervised prediction of inter-residue contacts [32]. Here, we introduce an even more direct strategy to open the black box of pLMs and infer coevolutionary information directly from the ESM-2 embedding space without supervised learning.

For a protein sequence of length *L*, the ESM-2 model generates an *L* × *M* (e.g., *M*=1280) embedding matrix. These embeddings have been widely used as inputs for downstream tasks such as structure prediction and protein functional annotation. Because the embedding captures the learned representation of sequence context, perturbing any residue in the input sequence alters the embedding in a manner that reflects the contextual importance of that residue. Here, we introduce perturbations by systematically masking the *i*-th residue and quantifying the resulting changes in the embedding (Figure 2a). The embedding difference projected onto the *j*-th residue, denoted *C_i_*( *j*), reports the contextual correlation strength between residues *i* and *j*, capturing coevolutionary dependencies encoded by the model (Methods). This mask-and-probe strategy provides an alternative to the traditional Multiple Sequence Alignment (MSA) based coevolutionary analysis in deriving coevolutionary signals from sequences [23–25]. The coevolutionary couplings derived from pLM embeddings provide biologically meaningful structural and functional information. As illustrated for two example proteins, i.e., the ribose-binding protein (RBP) and adenylate kinase (Adk) (Figure 2b), the coupling matrices inferred using this mask-and-probe based coevolutionary analysis closely mirror the distance maps from experimental structures [33, 34]. On a benchmark dataset [35], contact predictions from our mask-and-probe based coevolutionary analysis approach the accuracy of supervised neural-network predictors [13, 31, 36] and substantially outperform MSA based coevolutionary analysis methods [35] (Figure S1). These findings demonstrate a practical and interpretable strategy for directly decoding biologically relevant features from pLM embeddings.

**Figure 2.**
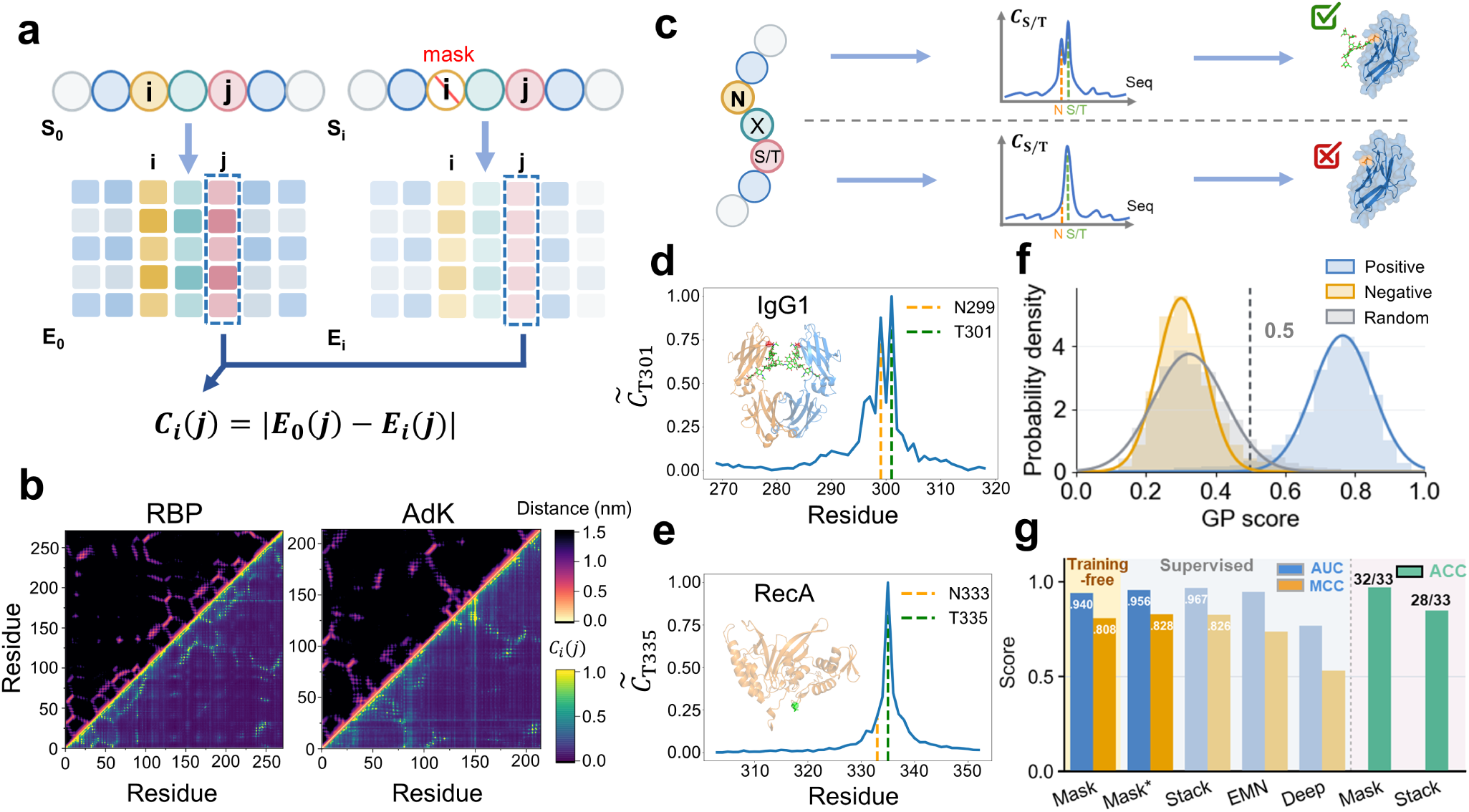
Deriving coevolutionary coupling and *N*-glycosylation propensity from pLM embeddings. (a) Schematic illustration of mask-and-probe strategy used to extract coevolutionary coupling profiles from ESM-2 embeddings. Masking residue *i* perturbs the embedding at residue *j*, and the resulting responses quantify the coupling strength between residues *i* and *j*. (b) Comparison between coevolutionary coupling matrices derived by the mask-and-probe strategy (lower) with the reference distance maps (upper) for two representative proteins RBP (PDB entry: 1URP [33]) and Adk (PDB entry: 1AKE [34]). (c) Overview of the MaskNGly workflow for predicting *N*-glycosylation sites from coevolutionary coupling profiles of canonical N-X-S/T sequons. (d,e) Representative coevolutionary coupling profiles for a glycosylated site, N299 of IgG1 (UniProt entry: P0DOX5), and a non-glycosylated site, N333 of *E. coli* RecA (UniProt entry: P0A7G6). (f) GP-score distributions for positive (blue), negative (yellow), and random (grey) datasets; solid lines denote Gaussian fits. (g) Comparison of training-free MaskNGly (Mask) with three supervised machine-learning predictors, including StackGlyEmbed (Stack)[19], EMNGly (EMN) [17], and DeepNGlyPred (Deep) [12], on the N-GlyDE test set. Mask^∗^ represents a minimal MLP classifier trained on a broader set of local coevolutionary coupling features derived from pLM embeddings. Right-hand bars: Prediction accuracies (Acc) on a smaller but more challenging test set of *N*-glycosylation sites with revised UniProt annotations. See also Tables S1-S4.

### Coevolution-based prediction of *N*-Glycosylation Sites

We next applied this mask-and-probe based coevolutionary analysis to investigate *N*-linked glycosylation sites, which occur in the canonical sequon N-X-S/T (where X̸=P). The task therefore is to distinguish the glycosylation-competent sequons from those that arise randomly. Because functional sequons are subject to evolutionary constraints, the Asn and Ser/Thr residues are expected to exhibit strong contextual coupling. Masking the Ser/Thr position should thus elicit a pronounced response at the corresponding Asn position in the embedding space (Figure 2c), and vice versa. As a demonstration, we examined the extensively characterized glycoprotein IgG1, which contains a conserved *N*-glycosylation site at N299 within the canonical sequon N299-X-T301 (Figure 2d). When T301 was masked, the resulting response curve displayed a prominent secondary peak at N299 in addition to the primary peak at the masked position. In general, the response strength *C_i_*( *j*) damps with increasing sequence separation between the mask site *i* and the probe site *j*. However, for authentic glycosylation sites, we observed a markedly elevated response at N299, denoted as *C*_T301_(N299) (Figure 2d), indicating strong coevolutionary coupling between N299 and T301. Conversely, masking N299 produced a reciprocal pattern: the response curve exhibited a strong signal not only at the masked N299 but also at the corresponding T301, quantified by *C*_N299_(T301) (Figure S2a). This bidirectional coupling effect reflects the functional interdependence of the two residues, as both sites are required to form a glycosylation-competent sequon. In contrast, non-functional N-X-S/T sequons (*i.e.*, those that will not be glycosylated) do not display such coupling as illustrated by a bacterial protein (RecA of *E. coli*) lacking *N*-glycosylation (Figure 2e, Figure S2b). These results demonstrate that the mask-and-probe based coevolutionary analysis effectively uncovers latent *N*-glycosylation signals encoded within pLM-embeddings.

To quantitatively characterize the coevolutionary coupling between the two residues in the canonical *N*-glycosylation sequon, we defined a glycosylation propensity (GP) score as *GP* = [*C̃*_S/T_(N) + *C̃*_N_(S*/*T)]*/*2. Here *C̃*_S/T_(N) represents the normalized *C*_S_*_/_*_T_(N), with N and S/T corresponding to the Asn and Ser/Thr sites of the *N*-glycosylation sequon (see Methods). We computed GP scores for all *N*-glycosylation sequons in a positive dataset and a negative dataset. The positive dataset consisted of ∼ 5826 experimentally verified *N*-glycosylation sites curated from UniProt [37], whereas the negative dataset comprised ∼ 11552 sequons curated in previous work from nuclear and mitochondrial proteins, which are known rarely *N*-glycosylated [9, 12, 38]. Strikingly, the distributions of the GP scores for the two datasets were clearly separated (Figure 2f and Figure S3). For a more robust test, the scores from the positive dataset were predominantly positioned above 0.5 with a Gaussian distribution, whereas those from the negative dataset were largely distributed below 0.5. A random dataset generated by selecting arbitrary residue pairs (*i* and *i* + 2, with *i* representing any sequence position) exhibited a distribution nearly identical to the negative dataset (Figure 2f). This similarity between the negative and random datasets indicates that, despite matching the canonical N-X-S/T sequon, negative sequons behave like random residue pairs with respect to glycosylation-related information, underscoring the specificity of the coevolutionary signal for biologically functional sequons.

The clear separation between the GP-score distributions of the positive and negative datasets indicates that the coevolutionary signals extracted from pLM embeddings can be directly used to predict *N*-glycosylation sites, yielding a training-free predictor of *N*-glycosylation sites (hereafter termed MaskNGly). To systematically assess its performance, we computed GP scores for the N-X-S/T sequons of proteins in the N-GlyDE test set curated previously from UniProt annotations [39], and classified each sequon as glycosylated or non-glycosylated. MaskNGly achieves an Area Under the Receiver Operating Characteristic Curve (AUC) and Matthews Correlation Coefficient (MCC) of 0.940 and 0.808, respectively, matching the performance of state-of-the-art supervised neural-network models (Figure 2g) [12, 16, 17, 19, 39]. The comparable performance suggests that the coevolutionary pattern extracted from ESM-2 embedding captures the essential sequence features governing N-glycosylation. A more detailed analysis further revealed an asymmetric correlation between the N and S/T positions: the response of the N site upon masking the S/T site is more strongly associated with glycosylation propensity than the reverse direction (Figure S3). The corresponding AUC score is slightly higher (0.947) (Table S1). Incorporating longer-range of coevolutionary couplings together with a multilayer perceptron (MLP) classifier yields only modest improvement (AUC=0.956, Methods), indicating the dominant role of the coevolutionary coupling between the N and S/T positions of the motif in decoding the *N*-glycosylation propensity.

The UniProt annotation is subject to continuous updates and corrections [37]. For example, the N-glycosylation dataset N-GlyDE, curated from the UniProt 2016_08 release and widely used to train and evaluate glycosylation-site predictors[12, 16, 17, 19, 39], contains both false-negative and potentially false-positive annotations when re-examined against the latest UniProt records (Table S3 and Table S4). These sites with corrected annotations therefore provide a more stringent benchmark for evaluating predictor performance. Among the sequons previously annotated as non-glycosylated in dataset N-GlyDE, 33 were updated as experimentally supported *N*-glycosylation sites based on the UniProt 2026_02 records, indicating false-negative labels in the original dataset. MaskNGly correctly predicted 32 of these revised positive sites. For these more challenging, annotation-revised sites, MaskNGly also achieves performance comparable to, or even higher than, that of supervised machine-learning models (Figure 2g). Conversely, 9 sites previously annotated as experimentally supported positives are no longer supported by updated and more stringent glycoproteomic evidence and are therefore treated as uncertain. For these sites, both MaskNGly and supervised models predict a significant portion of negative results (7/9 and 4/9, respectively), in line with the updates of UniProt annotations (Table S3 and Table S4). As such mis-annotations may be widespread in glycosylation datasets, they inevitably introduce biases into the predictions of supervised-learning models trained on them, underscoring the need for annotation-independent, training-free predictors.

### Experimental validation using independent glycoproteomic profiling

As an additional experimental validation of MaskNGly, we collected an independent set of glycoproteomic data using Click-iG, a recently reported chemical glycoproteomic strategy for intact glycopeptide-centric profiling [40, 41]. We applied Click-iG to human intrahepatic biliary epithelial cells (HIBEpiC), an *in vitro* model of the intrahepatic biliary epithelium (Figure 3a, see Methods). This experimental analysis identified 247 unique *N*-glycosylation sites across 146 glycoproteins. Notably, 111 of these sites (∼ 45%) were previously unreported and therefore absent from the UniProt knowledge base (Figure 3b). Most glycoproteins were detected with a single *N*-glycosylation site, whereas many sites showed glycan-composition microheterogeneity, being associated with multiple glycan composition types (Figure S4). We applied MaskNGly to all potential N-X-S/T sequons within these glycoproteins. The GP score distribution of experimentally detected sites showed a strong enrichment above the 0.5 threshold, closely mirroring the distribution of the positive reference dataset (Figure 3c). In contrast, negative and random datasets were largely restricted to GP scores below 0.5. Strikingly, more than 95% (106/111) of the newly identified positive sites received GP scores above this threshold, providing strong orthogonal validation that MaskNGly faithfully identifies functional glycosylation sites by directly reading out sequence-encoded signals from the pLM embedding, despite not applying any supervised training on annotated glycoproteomic data.

**Figure 3.**
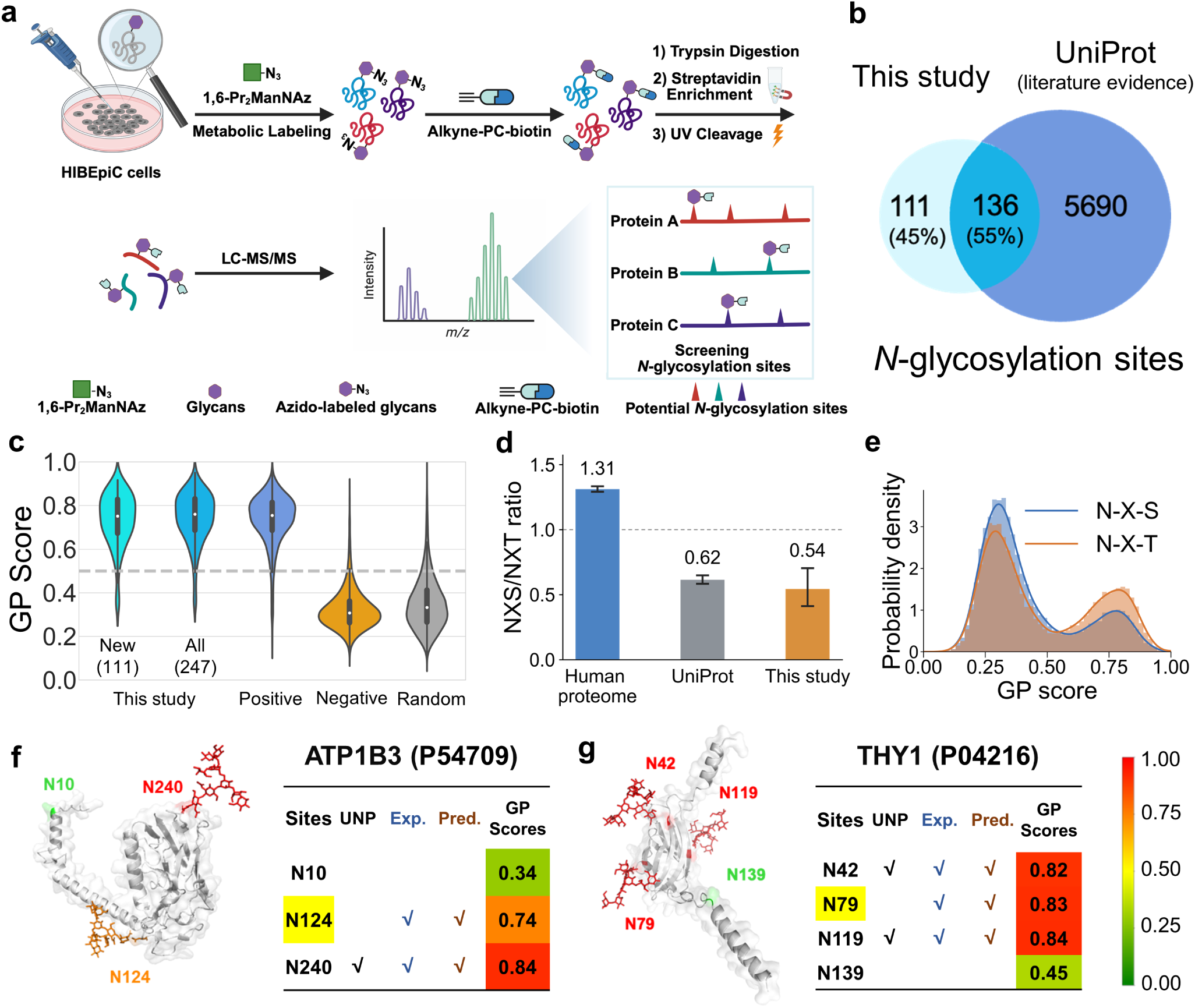
Experimental validation of MaskNGly predictions using intact glycoproteomics. (a) Schematic of the Click-iG strategy in HIBEpiC cells. HIBEpiC cells were metabolically labeled with 1,6-di-O-propionyl-N-azidoacetylmannosamine (1,6-Pr2ManNAz), reacted with alkyne–H-PC-biotin via click chemistry, digested and enriched by streptavidin beads, followed by photocleavage release of glycopeptides for whole glycopeptide analysis. (b) Venn diagram comparing the sizes of the datasets containing *N*-glycosylation sites identified in the experiment of this study with those annotated in the UniProt database. (c) Violin plots of MaskNGly predicted GP scores for test sets derived from the glycoproteomics data collected in this study, the positive dataset, the negative dataset, and the random dataset. For comparison, the GP score distribution for the newly identified sites is plotted separately. The dashed line marks the 0.5 threshold. (d) Ratios of N-X-S to N-X-T sequons in the human proteome, in UniProt annotations, and in the experimental dataset of this study. Error bars represent 95% site-level bootstrap confidence intervals computed from 100,000 resamples, and the dashed line denotes a ratio of 1. (e) Histograms and kernel-density curves of MaskNGly GP scores for human-proteome N-X-S and N-X-T sequons. (f, g) MaskNGly predictions for two representative glycoproteins, ATP1B3 (f) and THY1 (g). Left panels: cartoon representations of the protein structures (grey) with glycosylation sites and the modelled glycans colored according to the predicted GP score. Right panels: computed GP scores for each sequon, alongside the predicted glycosylation state (Pred.), UniProt annotation with experimental support (UNP), and glycosylation state identified in the experiment of this study (Exp.). A checkmark indicates a glycosylated site. GP scores are color-coded from green (low) to red (high).

Using these experimental annotations, we further quantified the relative frequencies of Ser and Thr at the third position of the N-X-S/T motif. Threonine was more frequently observed than serine among experimentally detected glycosylation sites (Figure 3d), consistent with the well-documented preference for N-X-T over N-X-S sequons in *N*-glycosylation [42–44]. The observed relative population of serine versus threonine was ∼0.54 (Figure 3d). A similar threonine enrichment was observed among experimentally supported UniProt *N*-glycosylation sites (Figure 3d). In contrast, threonine is less frequent than serine among all potential sequons in human proteins, with a serine versus threonine population of ∼1.31. These results indicate that N-X-S and N-X-T motifs have different glycosylation propensity, with N-X-T sites being more likely to undergo glycosylation. To assess whether this motif-level preference is captured by GP scores, we computed GP score distributions for all N-X-S and N-X-T sequons in human proteins. High-scoring sites were likewise enriched for N-X-T relative to N-X-S motifs (Figure 3e), demonstrating that GP scores can well capture this established sequence preference directly from protein pLM embeddings.

MaskNGly also predicted high GP scores for several sequons lacking prior experimental support but subsequently validated by our newly curated experimental dataset. For example, ATP1B3, which contains three canonical *N*-glycosylation sequons (N10, N124, N240), has experimental support only for glycosylation at N240, and MaskNGly correctly predicted a high GP score for this site (Figure 3f). Although N124 lacked prior annotation, MaskNGly predicted a high GP score, and our glycoproteomic experiment confirmed that this site is glycosylated. In contrast, the N10 sequon, which is unsupported by UniProt, received a low GP score (0.35), indicating low glycosylation propensity. For another protein THY1, MaskNGly predicted high GP scores for the sites N42, N79, and N119 (Figure 3g). Among these sites, N79 has not been experimentally validated previously but was identified as an *N*-glycosylation site in our experiment. Another site, N139, lacked experimental support and MaskNGly predicted a low GP score. Together, these results demonstrate that MaskNGly reliably translates sequence-level embeddings into accurate predictions of glycosylation propensity. Its ability to identify both known and previously unrecognized glycosylation sites highlights its value for guiding experimental discovery and underscores its broad utility in glycoproteomic studies.

### Proteome-scale decoding of *N*-glycosylation

#### Evolutionary divergence of N-glycosylation potential across domains of life

Because MaskNGly directly extracts glycosylation information from pLM embeddings and does not rely on annotated glycosylation datasets, it intrinsically has high transferability and can be applied to proteins from different species. We therefore extended high-throughput *N*-glycosylation site prediction to the proteome scale across domains of life. Using the AlphaFold Protein Structure Database (AlphaFold DB) [27, 45], we systematically analyzed all N-X-S/T sequons across the proteomes of diverse organisms (Figure 4a). Analysis of GP score distributions revealed that bacterial species dominantly exhibit unimodal distributions below 0.5, consistent with the near absence of

**Figure 4.**
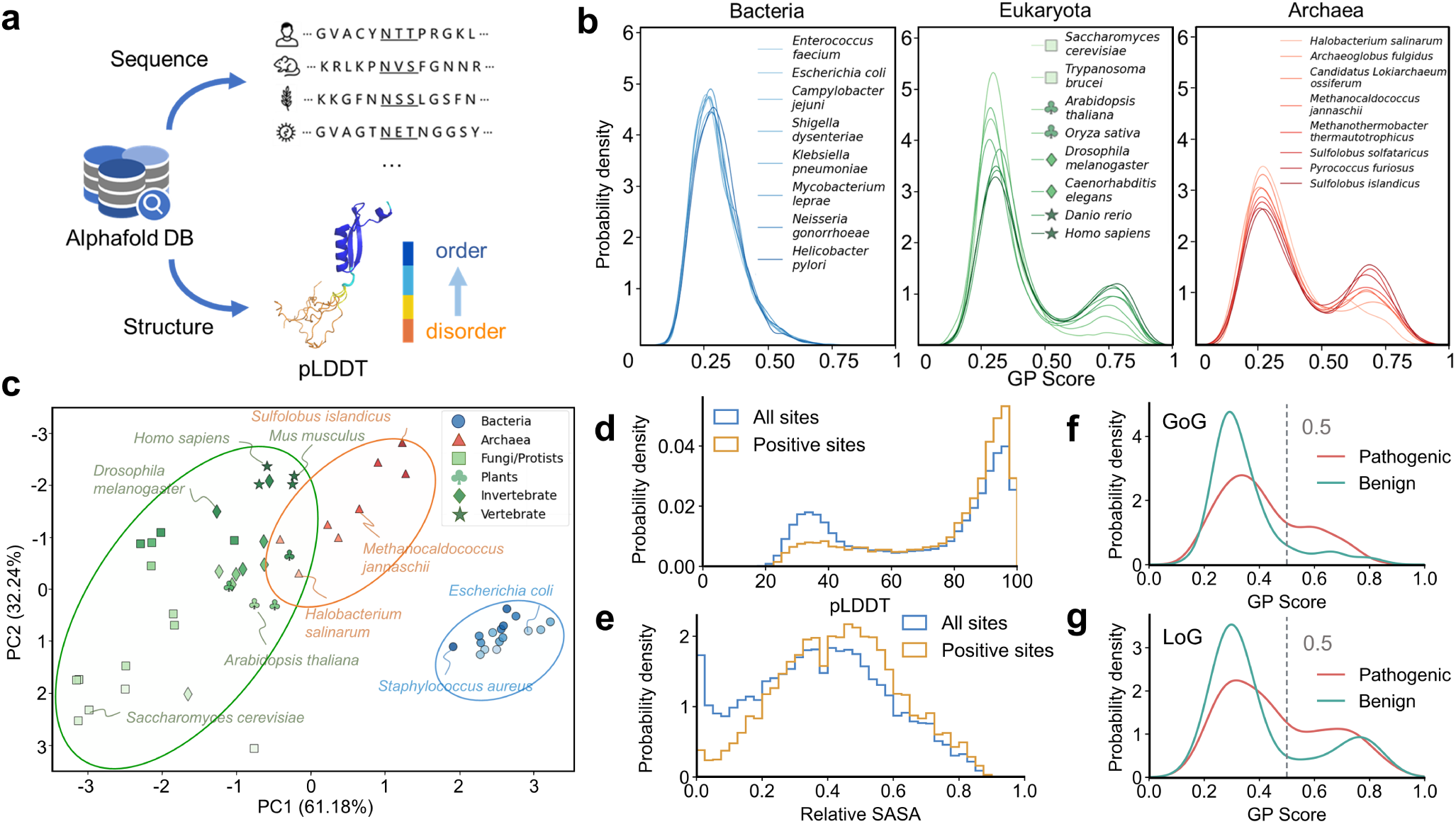
Proteome-scale decoding of *N*-glycosylation landscape. (a) Schematic illustrating the extraction of proteome-scale sequences and three-dimensional structures from AlphaFold database. (b) GP score distributions for all N-X-S/T sequons across proteins from Bacteria, Eukaryota, and Archaea. Curves are colored by mean GP score, with darker shades indicating higher values. (c) Principal-component analysis of GP-score distributions across species, showing separation among Bacteria, Eukaryota, and Archaea. Each distribution is represented by the fitted means and standard deviations of a two-component Gaussian mixture model in the principal-component analysis. (d, e) Distributions of pLDDT (d) and relative solvent-accessible surface area (rSASA) (e) for all candidate sequons (blue) and high GP-score sequons (GP *>* 0.5; yellow) in the human proteome. (f-g) GP-score distributions of the GoG (gain of glycosylation) and LoG (loss of glycosylation) variants. A total of 577 benign variants (GoG/LoG:284/293, cyan lines) and 519 pathogenic variants (GoG/LoG:311/208, red lines) are included.

*N*-glycosylation in bacterial proteins (Figure 4b). In contrast, eukaryotes, including simple unicellular species such as yeast, display a distinct secondary peak above 0.5, with this high-scoring population becoming progressively more prominent in more complex organisms ranging from unicellular organisms to vertebrates. Mean GP scores per organism further highlight this evolutionary divergence: bacteria predominantly show low average scores(∼ 0.30), whereas eukaryotes exhibit substantially higher values (0.34-0.46). Notably, both the mean scores and variance of these scores increased with biological complexity (Figure S5a), reflecting greater variability in glycosylation potential among eukaryotes. A similar trend was observed in the fraction of sites with scores exceeding 0.5 (Figure S5b). These findings align with established knowledge that *N*-glycosylation, while uncommon in bacteria, becomes progressively more prevalent and functionally diverse across eukaryotic evolution [46, 47].

Strikingly, archaeal species exhibit GP score distributions that more closely resemble those of eukaryotes than bacteria, displaying clear bimodal characteristics (Figure 4b). This observation is consistent with verified knowledge that archaea possess sophisticated *N*-glycosylation systems [48]. The experimental characterization of archaeal glycosylation remains challenging due to technical limitations [49], highlighting the value of our computational approach for exploring glycosylation in species that are difficult to study.

To further quantify these evolutionary patterns, we performed principal component analysis (PCA) on features extracted from the score distributions using a double-Gaussian mixture model (Methods). As shown in Figure 4c, the resulting PCA plot reveals clear separation among the three domains of life, with bacterial, archaeal, and eukaryotic clusters occupying distinct regions of the essential space. Intriguingly, within the Eukaryota cluster, we observe a pattern of continuous change that correlates with organismal complexity, likely reflecting the increasing sophistication of *N*-glycosylation machinery throughout eukaryotic evolution. This domain-specific clustering provides new insights into the origins and evolutionary diversification of protein glycosylation, suggesting that *N*-glycosylation signatures may serve as informative markers for studying protein evolution and species relationships.

#### Relationship between protein structure and N-glycosylation

*N*-glycosylation is an enzyme-catalyzed post-translational modification, and its occurrence is therefore expected to depend on the local structural features surrounding the modification site. Previous studies incorporating structural features into deep neural network models improved glycosylation predictions, suggesting the importance of protein structure in governing glycosylation modification [12, 18]. However, the structure-glycosylation relationship remains rather vague. The training-free nature of MaskNGly and the availability of proteome-scale structure data from AlphaFold DB make the extensive characterization of structure-glycosylation relationships possible. We next investigated the structural determinants that govern whether an N-X-S/T motif becomes glycosylated. Although it has long been proposed that stable and solvent-exposed regions of proteins are more conducive to glycosylation [10, 50, 51], a proteome-wide validation of these hypotheses has remained challenging. The computational framework developed here enables a systematic, cross-species examination of these structural factors.

We first investigated the relationship between protein structural integrity and glycosylation propensity. Structural integrity was quantified using predicted local distance difference test (pLDDT) scores from AlphaFold DB [45], which measures local confidence in the predicted structure. High pLDDT values correspond to well-folded regions, whereas low values indicate disordered conformations [52]. Intriguingly, the distribution of pLDDT for all potential *N*-glycosylation sites in human proteins exhibits a bimodal pattern (Figure 4d), indicating that sequons occur in both structured and disordered regions. In contrast, sites confidently predicted as glycosylated (score *>* 0.5) are predominantly located in well-structured regions (pLDDT *>* 80), showing a near-unimodal distribution pattern. Moreover, the percentage of positive sites increases steadily with pLDDT above a threshold of ∼ 30 (Figure S6a), supporting the notion that structural stability favors *N*-glycosylation. This trend is most pronounced in higher eukaryotes. Within the high-pLDDT region (pLDDT *>* 50), the percentage of positive sites per species increases markedly with organismal complexity, with bacteria consistently exhibiting low percentage of positive sites (Figure S7).

We next examined the effect of solvent accessibility on *N*-glycosylation propensity. To enable quantitative comparison, we defined the relative solvent-accessible surface area (rSASA) at residue-wise (Methods), which ranges from 0 (fully buried) to 1 (fully exposed). Potential sequons in human proteins span a broad range of solvent accessibilities, including buried regions (rSASA *<* 0.1) and solvent exposed regions (rSASA *>* 0.3) (Figure 4e). In contrast, positively predicted sites are strongly enriched in solvent-exposed regions and are rarely found in buried environments. The percentage of positive sites among potential sites increases with rSASA (Figure S6b), reflecting the requirement for surface accessibility during glycan transfer, as glycosylation processing machinery must be able to access the sequon. Cross-species comparisons reveal that the percentage of positive sites per species for solvent-exposed regions (rSASA *>* 0.3) increases markedly with increasing organismal complexity (Figure S8), mirroring the evolutionary pattern of the structure–glycosylation relationship.

#### Pathogenic consequences of sequon-altering mutations

To directly assess the functional importance of *N*-glycosylation, we examined mutations that alter glycosylation sequons and evaluated their association with disease. Mutations that create or disrupt *N*-glycosylation motifs, termed gain-of-glycosylation (GoG) and loss-of-glycosylation (LoG), respectively, can affect protein function and are therefore implicated in diverse pathologies [53, 54]. However, an omics-scale characterization of such correlation is still lacking. To systematically investigate these effects, we curated 1,096 single-point missense variants from the ProteinGym clinical variants substitutions database [55] that modify the canonical N-X-S/T motifs, comprising 595 GoG and 501 LoG variants annotated with pathogenicity.

It is worth noting that the above GoG and LoG variants are defined purely at the sequence level, and the altered sequons do not necessarily correspond to positively annotated glycosylation sites. This raises the question of whether the glycosylation propensity of these GoG and LoG sequons is related to pathogenic consequences. The GP score distributions of these altered motifs reveal striking patterns. For GoG variants, the majority of benign mutations introduce new sequons with low GP scores (*<*0.5) (Figure 4f). In contrast, a significantly higher proportion of pathogenic GoG mutations generate high-scoring new sequons (GP score *>*0.5), suggesting that acquisition of an *N*-glycosylation site through mutation more frequently leads to deleterious functional consequences. A similar trend is observed for LoG variants (Figure 4g). These results underscore the functional indispensability of naturally existing glycosylation sites for maintaining normal protein activity [3].

#### Characterizing glycosylation-mediated protein allostery

Beyond serving as static labels, glycans are increasingly recognized as dynamic regulators of protein conformation, stability, and long-range allosteric communication [56, 57]. A well-characterized example is human CD2, a glyco-protein that binds CD58 to trigger immune activation [58]. Experimental studies have shown that *N*-glycosylation at Asn65 is essential for binding with the downstream partner CD58 to activate immunity response, despite the glycan being spatially distant from the binding interface [59], implicating a long-range allosteric mechanism. Previous MD simulations further suggested that the Asn65 glycan forms specific hydrogen bonds with surface residues, including Asp16, Asn18, Asp20, Lys64, and Lys69, thereby stabilizing the CD2-CD58 complex through allosteric communication [57]. Such glycan-mediated allostery is expected to impose coevolutionary constraints on protein sequences, leaving detectable signatures in pLM embeddings. This raises the question of whether these allosteric signals can be extracted using our mask-and-probe based coevolutionary analysis.

Applying this approach to CD2, we first masked Asn65 and computed the resulting coevolutionary coupling profile, *C*_N65_( *j*), with *j* being the residue indices. The strongest responses were concentrated in two sequence regions, i.e., residues 14-26 and 58-72, which coincide with glycan-contacting regions identified by hydrogen-bond analysis of all-atom MD simulation trajectories (Figure 5a-c). This agreement indicates that our method can identify glycan-interacting residues from pLM embeddings, without structural input. We then performed the inverse analysis by masking each CD2 residue *j* and measuring the responses on Asn65 (Figure 5d). The resulting *C_j_*(N65) profile again highlighted the two glycan-contacting regions and additionally revealed an emergent peak at Trp35, a residue buried at the CD2-CD58 interface (Figure 5e). Trp35 lies adjacent to loop 1, a binding hotspot that has been implicated in relaying glycan-dependent perturbations in previous work [57], supporting its potential role in transmitting allosteric effects from Asn65 to the binding surface.

**Figure 5.**
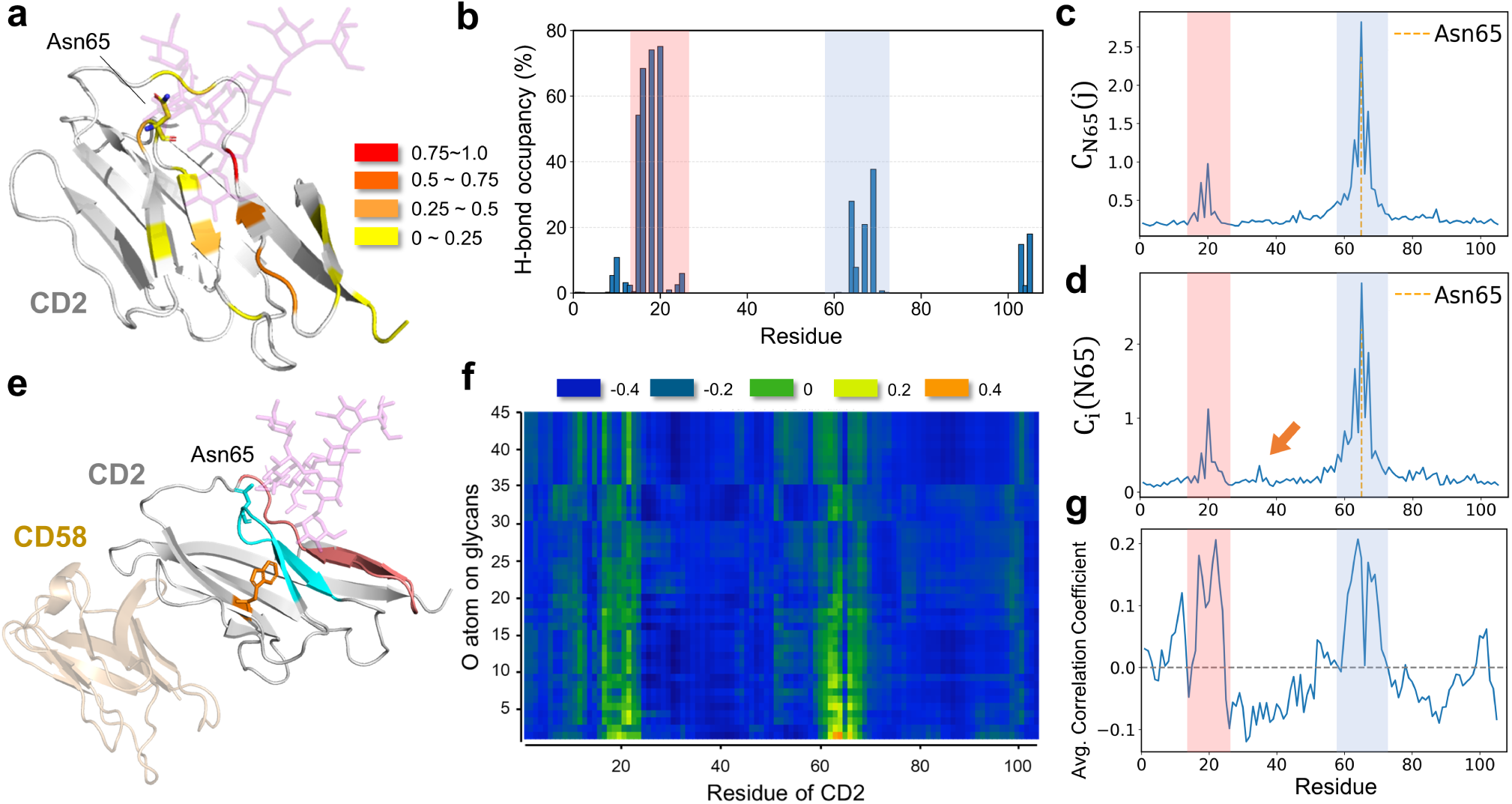
Inferring glycosylation-mediated allostery dynamics from pLM embedding. (a) Cartoon representation of glycosylated protein CD2 (PDB entry: 1GYA [61]). The glycan attached to Asn65 is shown as transparent purple sticks, and CD2 residues forming hydrogen bonds with the glycan are colored according to hydrogen-bonding occupancy from MD simulations, with red indicating higher occupancy. (b) Per-residue hydrogen-bonding occupancy between CD2 and the glycan derived from MD simulations. (c) Coevolutionary coupling profile induced by masking Asn65, *C*_N65_( *j*), revealing pronounced peaks within the highlighted regions. (d) Coevolutionary coupling profile, *C_i_*(N65), quantifying the responses of the embedding at Asn65 upon masking each residue *i*. The highlighted regions correspond to major coupling peaks, while the emergent peak at Trp35 (orange arrow) suggests a putative site mediating allosteric coupling between the CD2-CD58 interface and Asn65 glycosylation. (e) Cartoon representation of the CD2-CD58 complex (PDB entry: 1QA9 [58]). Trp35, buried at the binding interface, is shown in orange; CD58 is rendered semi-transparent for clarity. (f) Dynamic cross-correlation matrix (DCCM) between glycan oxygen atoms and CD2 C*α* atoms from MD simulations. (g) Per-residue average of the DCCM plotted along the CD2 sequence, with highlighted regions marking correlation peaks.

To further evaluate this embedding-derived allosteric pattern, we performed dynamic cross-correlation matrix (DCCM) analysis between glycan oxygen atoms and CD2 C*α* atoms based on all-atom MD simulations (Figure 5f-g) [60]. The residue-wise correlation profile showed positive correlation peaks within the same two regions as those identified by the mask-and-probe based coevolutionary coupling profiles. In contrast, Trp35 was located in a local negative-correlation valley, consistent with its role as a distal interface residue dynamically coupled to glycan motion rather than directly contacting the glycan. Thus, the DCCM results independently support the spatial and dynamic relevance of the coupling regions inferred from pLM embeddings.

The strong concordance between our embedding-based predictions and MD simulations demonstrates that the mask-and-probe strategy captures functionally relevant, dynamic features of glycans on glycoproteins, indicating that glycan-mediated allostery leaves detectable evolutionary signatures in pLM embeddings. By identifying these evolutionary correlations directly from pLM embeddings without supervised training, our method provides a rapid, scalable alternative to MD simulation-based approaches for probing glycan-dependent allosteric mechanisms.

## Discussion

Protein glycosylation mediates diverse biological processes and is tightly regulated by cellular context [3–5]. Throughout evolution, protein sequences have adapted not only to optimize intrinsic structure and function but also to coevolve with post-translational modification systems [51]. Decoding glycosylation information directly from sequence therefore remains a fundamental yet challenging problem. This work demonstrates that protein language model embeddings contain a directly accessible signature of *N*-linked glycosylation for a given N-X-S/T sequon. Building on this observation, we developed MaskNGly, a training-free and interpretable framework that decodes *N*-glycosylation signals directly from pLM embeddings using a mask-and-probe based coevolutionary analysis. By systematically perturbing sequence inputs and quantifying embedding responses, MaskNGly uncovers residue–residue couplings that recapitulate functional sequons and predict glycosylation propensity at the proteome scale. Remarkably, this approach achieves performance comparable to state-of-the-art supervised machine-learning predictors without reliance on annotated training sets.

Proteome-scale application of MaskNGly revealed marked differences in the distribution of sequence-encoded glycosylation propensity across the three domains of life. Bacterial proteomes were dominated by low-scoring sequons, whereas archaeal and eukaryotic proteomes contained a distinct high-scoring population. Among eukaryotes, both the mean GP score and the fraction of high-scoring sequons tend to increase with organismal complexity, indicating substantial cross-species variation in the sequence-encoded potential for *N*-glycosylation. In the context of eukaryotic glycosylation-system evolution, this trend is particularly noteworthy. Cross-species glycoproteomic studies have shown that, although the core biosynthetic machinery of *N*-glycosylation is broadly conserved, the repertoire of glycoprotein substrates has diversified markedly across lineages[62]. Glycan-processing pathways likewise exhibit pronounced phylogenetic specificity: yeast predominantly generates high-mannose structures [63]; the Drosophila glycome remains enriched for high-mannose and paucimannose species with limited hybrid and complex forms[64]; and vertebrates, including humans, possess highly elaborated processing networks capable of extensive branching, terminal modification, and structural heterogeneity[65, 66]. These observations illustrate that glycosylation diversification is typically manifested through the abundance and heterogeneity of glycoforms among different species. The present results instead demonstrate a positive correlation between organismal complexity and the intrinsic glycosylation propensity of sequons themselves. This suggests that the sequence-encoded potential for glycosylation may have expanded in parallel with lineage-specific elaboration of glycan-processing capacity, thereby broadening the regulatory space accessible to glycan-mediated molecular control. Together, these findings provide a sequence-level perspective on how glycosylation capacity has been shaped across species evolution.

The structural analysis provides a complementary characterization of the biophysical basis underlying the glycosylation landscape. Across the human proteome, using AlphaFold2-predicted structures and confidence scores (pLDDT), we find that high-scoring human sequons were enriched at residues with high AlphaFold pLDDT and greater solvent accessibility, consistent with the requirement that glycosylation sites be presented in accessible local contexts [10, 50]. The depletion of high-scoring sites from low-confidence and therefore often disordered regions suggests that functional sequons are preferentially embedded in locally structured contexts. These observations may imply that glycosylation processing machinery utilizes not only sequence motif but also local structural features in recognizing specific glycosylation sites. Interestingly, recent studies show that incorporating structural information alongside sequence features enables supervised deep-learning models to achieve improved prediction of *N*-glycosylation sites, consistent with the proteome-scale correlation between local structural features and glycosylation propensity revealed in this work [18, 20].

The variant analysis of glycosylation propensity using the ProteinGym clinical variants substitutions database further extends the framework from site prediction to the assessment of pathogenic consequences. Perturbations to motif glycosylation propensity correlate tightly with pathogenic outcomes: GoG and LoG variants were more often pathogenic when they created or disrupted high-scoring sequons, respectively. This strong association between glycosylation motif-altering mutations and pathogenicity underscores the importance of incorporating glycosylation potential into genetic variant interpretation. We also showed that coevolutionary analysis of pLM embeddings can be used to identify residue networks potentially involved in glycan-mediated allostery, which otherwise requires computationally expensive MD simulations. The close agreement between embedding-derived couplings and MD-based hydrogen-bonding and dynamic cross-correlation profiles further supports the biological relevance of these coevolutionary couplings from pLM embeddings.

Several limitations also highlight opportunities for future development. The current method is restricted to the canonical N-X-S/T sequons. Decoding modifications that lack a universal consensus sequon, such as nucleocytoplasmic O-GlcNAcylation and mucin-type O-GalNAc glycosylation [65, 67], will require more sophisticated inference strategies and more comprehensive experimental benchmarks. Incorporating pLM embeddings not only from glycoproteins but also from the proteins involved in the glycosylation processing pathways may further constrain sequence features governing *O*-glycosylation propensity. In addition, this work focuses primarily on the accurate prediction of glycosylation sites. However, the efficient and precise inference of glycoforms and glycan conformations represents equally important but substantially more challenging tasks owing to the scarcity of high quality experimental data [21]. A promising direction for future research is the integration of pLM embeddings with emerging glycomics measurements and biophysical constraints, enabling a unified framework capable of decoding both glycoforms and three-dimensional glycan architecture directly from protein sequences. It is worth emphasizing that the current computational strategy is interpretable primarily in the sense that it directly infers biologically relevant coevolutionary statistics from pLMs without relying on supervised training. A more promising yet challenging direction for future work is to more closely relate these statistical signals to the underlying biophysical principles governing protein glycosylation, thereby moving toward fully opening the black box of pLMs and revealing how sequence-encoded constraints shape glycosylation across evolution.

Despite these caveats, the ability to probe glycan-dependent relationships directly from sequence embeddings, independent of curated annotations or explicit structural models, opens new avenues for studying post-translational modification biology at omics-scale. By bridging evolutionary signal, structural context, and functional consequence, embedding-based decoding offers a promising route toward deciphering the latent “glyco-code” and advancing glycosylation-based applications in biology and medicine. Moreover, accurate and efficient decoding of glycosylation profiles from pLM embeddings provides a foundation for the reverse problem, i.e., glycoprotein design. Introducing new glycosylation motifs into proteins could enable more flexible regulation of biological processes and facilitate the development of more effective protein therapeutics [68, 69]. Mask-and-probe based coevolutionary analysis of pLM embeddings thus holds substantial promise for both mechanistic discovery and rational engineering of glycoproteins.

## Methods

### Deriving coevolutionary couplings from pLM embeddings

For a given sequence, a perturbed sequence was generated by replacing the residue at position *i* with a `<mask>` token while leaving all other positions unchanged. Both the original sequence (**S**_0_) and masked sequences (**S***_m_*) were independently processed by ESM-2 (model: esm2_t33_650M_UR50D) to obtain their corresponding embedding matrices, **E**_0_ and **E***_m_*. Each embedding matrix has dimensions *L* × 1280, where *L* denotes the sequence length. A difference matrix **E***_m_* − **E**_0_ was computed, and the L2 norm was applied along the embedding dimension to yield a response vector **C***_i_*( *j*) of length *L*, with *i* and *j* ranging from 1 to *L*. Here **C***_i_*( *j*) quantifies the response of the sequence embedding at position *j* when position *i* is masked, reflecting the coevolutionary coupling between the two positions. Strong functional coupling between residues *i* and *j* is expected to produce a large **C***_i_*( *j*). Similarly, masking position *j* and probing position *i* yields **C** *_j_*(*i*).

### Quantifying glycosylation propensity

For the canonical N-X-S/T sequon, functional *N*-glycosylation implies evolutionary coupling between the Asn and Ser/Thr positions. Masking the Asn position produces a strong embedding response at the Ser/Thr, and masking the Ser/Thr position produces a reciprocal response at the Asn position. In contrast, non-functional or randomly occurring sequons exhibit weak responses. To quantify glycosylation propensity, we define *GP* = (*C̃*_N_(S*/*T) + *C̃*_S/T_(N))/2. Here *C̃_i_*(*j*) denotes the min-max normalized value of *C_i_*( *j*). Thus, *GP*_N_ = *C̃*_N_(S*/*T) represents the response at the S/T position when masking N, and *GP*_S_*_/_*_T_ = *C̃*_S/T_(N) represents the reciprocal response at N when masking S/T. A threshold of 0.5 was applied to classify sequons as glycosylated or non-glycosylated, based on score distributions from independent test datasets (Figure 2f). For convenient access, the computational framework MaskNGly has been deployed as a public web server (https://maskngly.com).

### Inferring contact map from pLM embedding

As a demonstration that coevolutionary coupling extracted from pLM embedding contain meaningful biophysical information, we used the coupling profile *C_i_*( *j*) derived from the mask-and-probe coevolutionary analysis to predict protein contact maps. This approach leverages the well-established observation that spatially contacting residues are frequently evolutionarily coupled. Performance was assessed on the PSICOV150 benchmark dataset [35], comparing contact predictions obtained from pLM-inferred *C_i_*( *j*), an MSA-based coevolutionary contact predictor PSICOV [35], the contact prediction module of ESM-2, which infers residue contacts from attention-based representations by using trained logistic regression[13, 31], and a deep convolutional neural network model DeepCov [36] (Figure S1). The PSICOV150 benchmark dataset comprises 150 single-domain, monomeric proteins. The protein structure with PDB ID 1vjk was excluded due to inconsistent lengths of sequence and structure data, yielding 149 proteins for evaluation. Reference contact maps were generated from experimentally determined structures. Two residues were defined as contacting if their *C_β_* atoms were within 8 Å, with *C_α_* used for glycine. Following the PSICOV150 benchmark, residue pairs with |*i* − *j*| *>* 6 were evaluated as all contacts, whereas those with |*i* − *j*| *>* 23 were evaluated as long-range contacts.

To convert coevolutionary coupling profile into contact predictions, *C_i_*( *j*) score matrix was first symmetrized ([*C_i_*( *j*) + *C_j_*(*i*)]*/*2) and then subjected to average product correction to reduce residue-specific background effects [70]. The corrected coupling scores were subsequently standardized to zero mean and unit variance for the considered contact collections of each protein. ESM2 contact scores were produced by the contact prediction module of ESM-2 (model: esm2_t33_650M_UR50D). DeepCov predictions were generated from the official PSICOV150 alignments using the released DeepCov v1.0 covariance model with a 41-residue receptive field and the released feature-generation code and parameters [36]. PSICOV scores were obtained from the original predictions distributed with the PSICOV150 benchmark [35]. The corresponding areas under the receiver operating characteristic curve (AUCs) were calculated based on these scores and reference contact labels.

### Datasets

To evaluate the performance of MaskNGly, we used three test datasets: a positive dataset, a negative dataset, and a random control dataset (Figure 2f). The positive dataset was constructed by extracting experimentally verified *N*-linked glycosylation sites from UniProt, retaining only those sequons that satisfied the canonical N-X-S/T motif (X ≠ P) and were supported by at least one published experimental study (ECO:0000269) [37]. This resulted in a total of 5,826 high-confidence glycosylation sites. Although this dataset is relatively small compared to the whole UniProt dataset with glycosylation annotation, the requirement for direct experimental validation provides a more stringent and reliable benchmark for model evaluation. Because MaskNGly operates without supervised training, this dataset size is fully adequate for performance assessment purposes. In addition, we curated an independent experimental dataset derived from human intrahepatic biliary epithelial cells (HIBEpiC) using intact glycopeptide analysis as described below. All identified sites conformed to the N-X-S/T motif, resulting in 247 positive glycosylation sites with high-confidence mass spectrometry support.

The negative dataset was adopted from DeepNGlyPred [12] and consists of 11,552 non-glycosylated N-X-S/T motifs located in nuclear or mitochondrial proteins, as annotated by DeepLoc-1.0 [38]. These subcellular localizations are considered unlikely to support *N*-glycosylation [9], thereby providing high-confidence negative examples. Finally, a random control dataset was generated by randomly sampling an equal number of sites (5,826) from the same set of glycoproteins used in the positive dataset, without imposing the motif requirement. This dataset serves as a control to compare score distributions against biologically relevant sites.

### Evaluation metrics

For comparison with published predictors, MaskNGly was additionally evaluated on the standardized N-GlyDE test dataset (446 sites from UniProt release 2016_08) widely used in previous studies [12, 16, 17, 19, 37]. Performance was assessed using nine commonly applied classification metrics: Sensitivity (Recall), Specificity, Balanced Accuracy, Accuracy, Precision, F1-score, Matthews Correlation Coefficient (MCC), the Area Under the Receiver Operating Characteristic Curve (AUC), and the Area Under the Precision-Recall Curve (AUPR).

### Supplementary benchmark

To further assess whether the coevolutionary coupling between the N and S/T positions of the N-X-S/T motif extracted by the mask-and-probe strategy reflects the essential feature determining *N*-glycosylation site, we also compared MaskNGly with a multilayer perceptron (MLP) classifier trained on a broad set of local coupling signals around the motif. The MLP was trained using the same training dataset of *N*-glycosylation sites used in previous studies [12, 16, 17, 19, 39]. For each glycosylation site in the training set, we computed *C̃*_N_(*i*) and *C̃*_S/T_(*i*) with *i* ranges among the five sites flanking the center Asn of the canonical motif. This yielded a 10-dimensional input feature vector capturing a broader local coupling pattern around the canonical sequon. The MLP consisted of 10 input units, a single hidden layer with 100 ReLU-activated neurons and a sigmoid output unit. Training was performed using Adam with a learning rate of 0.001 and an L2 regularization of 0.0001. Sites with predicted positive-class probabilities greater than 0.5 were classified as glycosylated. Model performance was evaluated on the independent N-GlyDE test set [39], enabling direct comparison between MaskNGly and supervised learning approaches that incorporate wider contextual coupling information.

### Proteome-scale analysis

We downloaded AlphaFold2–predicted proteomes for 48 organisms from the AlphaFold Protein Structure Database [27, 45]. Proteomes for seven additional archaeal species were obtained from NCBI Datasets [71]. Per-residue pLDDT scores were extracted from the B-factor field of the corresponding PDB files. Solvent-accessible surface area (SASA) profiles were computed using MDTraj [72] with the Shrake–Rupley algorithm [73]. The resulting SASA values were then normalized by the maximal SASA of asparagine (1.95) to obtain the relative solvent-accessible surface area (rSASA) for each residue. To characterize the pathogenic consequences of N-glycosylation, we used the clinical variants substitutions database from ProteinGym [55]. From this resource, we curated 1,096 single-point missense variants that modify the canonical N-X-S/T motifs. Among these, 595 variants introduce a new canonical sequon (GoG, gain-of-glycosylation), whereas 501 variants disrupt an existing sequon (LoG, loss-of-glycosylation). The PCA analysis for the GP-score distributions across species was performed based on the fitted means and standard deviations with a two-component Gaussian mixture model.

### Experimental Validation

#### Cell Lines and Cell Culture

HIBEpiC cells were purchased from Shanghai Zhong Qiao Xin Zhou Biotechnology (Shanghai, China). HIBEpiC cells were identified using short tandem repeat profiling and tested negative for mycoplasma contamination. HIBEpiC cells were cultured in the epithelial cell medium supplemented with 1% epithelial cell growth supplement (EpiCGS), which was purchased from Shanghai Zhong Qiao Xin Zhou Biotechnology (Shanghai, China). Cells were cultured in a cell incubator at 37 ^◦^C with 5% CO2.

#### Reagents

Dimethyl sulfoxide (DMSO, cat. no. D2650), CuSO4 (cat. no. 931071), sodium ascorbate (cat. no. A7631), methanol (cat. no. 439193), urea (cat. no. U5378), ammonium bicarbonate (ABC, cat. no. A6141), dithiothreitol (DTT, cat. no. D9779), iodoacetamide (IAA, cat. no. I1149) were purchased from Sigma-Aldrich. Alkyne-PC-biotin (catalog no. BB-28), BTTAA (catalog no. BDJ-4), were purchased from Confluore Biotech (Xi’an, China). Alkyne-Cy5 (catalog no. TA116) was obtained from Click Chemistry Tools (Scottsdale, AZ, USA). 1,6-Pr2ManNAz was gifted from Dr. Ran Xie’s lab. Pierce bicinchoninic acid (BCA) Kit (cat. No. 23227) and Hoechst 33342 (cat. no. H3570) were purchased from Thermo Fisher Scientific. Mass spectrometry grade trypsin (catalog no. HLS TRY001C) was purchased from Beijing Life Proteomic. All of the organic solvents were of analytic grade.

#### Metabolic Oligosaccharide Engineering (MOE) of Living Cells

HIBEpiC cells seeded at 10-cm dishes at approximately 30% confluence were treated with 200 *µ*M 1,6-di-O-propionyl-N-azidoacetylmannosamine (1,6-Pr2ManNAz) for 48 h. The cells were harvested by trypsin digestion and washed twice with PBS and were lysed as described in the Protein Extraction. All procedures were performed under reduced light exposure to protect the photocleavable linker.

#### Cell lysis and protein extraction

The cells were harvested and re-suspended in PBS (pH 7.4, Gibco) or 4% Sodium dodecyl sulfate (SDS, wt/vol) in Milli-Q water. After lysing with sonication, the supernatant was collected. The pellet from the PBS group was re-suspended in 4% SDS and lysed again, of which the supernatant was also collected. After determination of the protein concentration using the BCA protein assay kit (Pierce), the cell lysates containing the same concentrations of proteins and SDS in PBS were obtained for further use.

#### Glycopeptide enrichment by Click-iG

The lysates extracted from HIBEpiC cells were incubated with 500 *µ*M premixed CuSO4/BTTAA (molar ratio 1:2), 100 *µ*M alkyne-photocleavable linker-biotin (alkyne-PC-biotin), and 2.5 mM freshly prepared sodium ascorbate in PBS (pH 7.4) for 3 h at room temperature. The mixture was added to 8 volumes of ice-cold methanol for precipitation overnight at −80 ^◦^C and washed three times with ice-cold methanol. Then, the protein pellet was reconstituted by sonication using 4 M urea in 50 mM ammonium bicarbonate (ABC), incubated with 10 mM DTT at 37 ^◦^C for 1 h, and followed by incubation with 20 mM iodoacetamide (IAA) at room temperature for 30 min in the dark. The solution was diluted to 1 M urea in 50 mM ABC supplemented with mass spectra grade trypsin (enzyme: substrate ratio at 1:50, w/w) and reacted at 37 ^◦^C for 16 h. After that, the streptavidin agarose beads (150 *µ*L per 40 mg, Thermo Fisher Scientific, 20359) were added to the solution above and gently rotated for 3 h at room temperature. The beads were washed five times with PBS and Milli-Q water successively and resuspended with 200 *µ*L 0.1% formic acid (FA, v/v), followed by irradiating three times under 365 nm UV light for 5 min using a UV cross-linker (CL-1000; UVP). The supernatant was collected, evaporated in a vacuum centrifuge, and subjected to LC–MS/MS analysis.

#### LC-MS/MS Analysis and Data Processing

The enriched glycopeptides were dissolved in 0.1% FA and subjected to LC-MS/MS analysis using an Orbitrap Fusion Lumos Tribrid mass spectrometer (Thermo Fisher) interfaced with a Nanospray Flex ion source and an EASY-nLC 1200 nanoflow UHPLC system. Full-scan MS spectra were recorded across m/z 350-2000 at a resolving power of 120,000 (at m/z 200), with monoisotopic precursor selection enabled. For MS/MS acquisition, multiply charged ions (z = 2-8) were isolated using a 3-s top-speed duty cycle, 15-s dynamic exclusion, and a ±10 ppm isolation window. Fragmentation was carried out using the sceHCD-pd-EThcD routine: HCD with a fixed first mass of m/z 100, NCE of 30 ± 10%, and resolution of 30,000 (at m/z 200), followed by product-dependent EThcD triggered upon detection of diagnostic oxonium ions (e.g., m/z 168.065, 186.076, 204.087, 274.092, 292.103, 300.130, 329.146, 342.178, 366.140, 388.146, 399.199, and 405.213; mass tolerance ±10 ppm) in the HCD spectrum. The supplemental activation energy for EThcD was set to 35.

Raw data were processed with pGlyco3 in the “HCD+EThcD” mode. The protein database was the SwissProt Homo sapiens proteome (UniProt, downloaded on 2016-11-04). N-glycan identification was performed against the built-in pGlyco-N-Human glycan database with glycan legality checking enabled. Precursor and fragment mass tolerances were set to ±10 ppm and ±20 ppm, respectively, with all other parameters kept at default. For samples labeled with ManNAz-derived probes, SiaNAz-H_2_O (m/z 387.13902, defined as PM) was included as a variable glycan modification substituting Sia (A)-H_2_O.

### Molecular dynamics simulations

The initial structure of the human CD2-CD58 complex was prepared from the crystal structure of HsCD2ad-CD58 (PDB entry: 1QA9 [58]). The glycan attached to Asn65 was modeled according to the glycosylated CD2 structure (PDB entry: 1GYA [61]). Missing residues and point mutations in the crystallographic complex were repaired following the previously reported protocol [57]. MD simulations were performed using GROMACS 2021.2 [74]. The system was solvated in a cubic TIP3P water box and neutralized with counterions [75]. Periodic boundary conditions were applied, and long-range electrostatic interactions were treated using the particle mesh Ewald method [76]. The Amber14SB force field was used for the protein, and the GLYCAM06-j force field was used for the glycan [77, 78]. We prepared three independent CD2-CD58 complex systems. These systems were first energy minimized and then heated to 300 K and relaxed for 0.1 ns under NVT ensemble, which was further followed by another round of relaxation simulations under NPT ensemble at 300 K and 1 atm for 0.1 ns. During the above relaxation simulations, a harmonic positional restraint was applied to the heavy atoms of the system with a spring constant of 1000 kJ/mol/nm^2^. Then the production simulations were conducted under NPT ensemble at 300 K and 1 atm for 1000 ns with the time step of 2 fs. Hydrogen-bond and DCCM analysis were carried out using MDTraj [60, 72] based on three independent trajectories.

## Acknowledgements

This work was supported by Basic Research Program of Jiangsu Province (BK20253050), National Natural Science Foundation of China (Grant Nos. 12574224, 12347102, 22477005 and 92478107), the Fundamental and Interdisciplinary Disciplines Breakthrough Plan of the Ministry of Education of China (JYB2025XDXM502), the Young Talent Program, SCF of Jiangsu Province, the Open Project Program of the State Key Laboratory of Medical Proteomics (SKLP-O202508) and the grant of Wenzhou Institute, University of Chinese Academy of Sciences (WIUCASQD2021010, WIUCASQD2022036). The authors also acknowledge support from Nanjing Kunpeng&Ascend Center of Cultivation, the HPC Center of Nanjing University, e-Science center of Nanjing University, and the Nanjing Key Laboratory for Cardiovascular Information and Health Engineering Medicine (funded by the Nanjing Municipal Health Commission) and its Jiangsu counterpart.

## Data availability

The data supporting this study, including the experimental glycoproteomic data generated in this work, are available at https://box.nju.edu.cn/d/7aa0a97985b3492eaa09. Our web server at https://maskngly.com provides online prediction of *N*-glycosylation sites by MaskNGly from user-submitted protein sequences, as well as queries of deposited across-species *N*-glycosylation predictions using UniProt accessions. The MaskNGly software package is available at https://github.com/ccbry/maskngly. The MD simulation data have been deposited at https://scidc.nju.edu.cn/records/b5hj8-2bd18.

## Author contributions

X.Z. and W.L. conceived the project. X.Z., X.G., and W.L. designed the algorithms. X.Z. and X.G. performed the computational analysis. X.Z., W.R. and Z.P. prepared the databases, software scripts, and web server. Y.Z., J.W., and R.X. conducted the glycoproteomic experiment. X.Z. conducted MD simulations. X.Z., W.L., R.X., and W.W. wrote the manuscript with input from all authors. W.L., R.X., and W.W. supervised the research.

