## Supplementary Tables and Figures for "Omics-scale decoding of protein *N*-glycosylation using protein language model embeddings"

**Table S1.** Performance comparison of MaskNGly with existing supervised machine-learning predictors on the N-GlyDE dataset. Double dashes indicate values that were not reported in the corresponding publications.

| Model | Sensitivity | Specificity | Balanced Accuracy | Accuracy | Precision | F1-score | MCC | AUC | AUPR |
|---|---|---|---|---|---|---|---|---|---|
| MaskNGly (average) | 0.935 | 0.888 | 0.911 | 0.906 | 0.836 | 0.883 | 0.808 | 0.940 | 0.854 |
| MaskNGly (mask N) | 0.941 | 0.841 | 0.891 | 0.879 | 0.783 | 0.855 | 0.762 | 0.936 | 0.858 |
| MaskNGly (mask S/T) | 0.959 | 0.852 | 0.905 | 0.892 | 0.798 | 0.871 | 0.790 | 0.947 | 0.880 |
| MaskNGly (trained) | 0.953 | 0.892 | 0.922 | 0.915 | 0.843 | 0.894 | 0.828 | 0.956 | 0.895 |
| StackGlyEmbed[a] [1] | 0.982 | 0.868 | 0.925 | 0.910 | 0.815 | 0.891 | 0.826 | 0.967 | 0.931 |
| LMNglyPred[a] [1, 2] | 0.747 | 0.942 | 0.845 | 0.869 | 0.886 | 0.810 | 0.718 | 0.845 | 0.756 |
| DeepNGlyPred[a] [1, 3] | 0.725 | 0.811 | 0.768 | 0.779 | 0.695 | 0.710 | 0.531 | 0.768 | 0.607 |
| EMNGly[b] [4] | 0.975 | 0.700 | – | 0.884 | – | – | 0.736 | 0.946 | – |
| N-GlyDE[c] [5] | 0.826 | 0.689 | – | 0.740 | 0.613 | – | 0.499 | – | – |
| GlycoMine[c] [5, 6] | 0.700 | 0.739 | – | 0.725 | 0.616 | – | 0.430 | – | – |

In the evaluation of MaskNGly using the N-GlyDE dataset, updates in UniProt annotations were considered.
[a] Performance values were taken from Ref. [1].
[b] Performance values were taken from Ref. [4].
[c] Performance values were taken from Ref. [5].

**Table S2.** Methodological comparison of MaskNGly and existing supervised machine-learning predictors. The training-free MaskNGly variants perform direct prediction from ESM-2 readouts without requiring task-specific supervised training or protein structural information.

| Model | Training-free | Structure required | Main representation |
|---|---|---|---|
| MaskNGly (average) | **Yes** | No | ESM2 direct readout; $GP = [\widetilde{C}_{\mathrm{N}}(\mathrm{S/T}) + \widetilde{C}_{\mathrm{S/T}}(\mathrm{N})]/2$ |
| MaskNGly (mask N) | **Yes** | No | ESM2 direct readout; $GP_{\mathrm{N}} = \widetilde{C}_{\mathrm{N}}(\mathrm{S/T})$ |
| MaskNGly (mask S/T) | **Yes** | No | ESM2 direct readout; $GP_{\mathrm{S/T}} = \widetilde{C}_{\mathrm{S/T}}(\mathrm{N})$ |
| MaskNGly (trained) | No | No | MLP using only $\widetilde{C}_{\mathrm{N}}(i \pm 2)$ and $\widetilde{C}_{\mathrm{S/T}}(i \pm 2)$ as 10-dimensional input features, i means the position of N. |
| StackGlyEmbed | No | No | ESM2, ProteinBERT, and ProtT5 embeddings combined using a stacking ensemble |
| LMNglyPred | No | No | ProtT5 embeddings with a supervised classifier |
| DeepNGlyPred | No | No | Supervised classifier |
| EMNGly | No | Yes | ESM-1b and AlphaFold2-derived features with a supervised classifier |
| N-GlyDE | No | No | Supervised classifier |
| GlycoMine | No | No | Supervised classifier |

**Table S3.** Results of MaskNGly predictions of *N*-glycosylation sites for a false-negative dataset with revised annotations. All sites were previously annotated as negative but were subsequently revised to positive. MaskNGly predictions were assigned using a score threshold of 0.5. "Previous" and "Revised" denote annotations based on UniProt releases 2016_08 and 2026_02, respectively.

| UID | Site | Previous | Revised | MaskNGly score | Prediction |
|---|---|---|---|---|---|
| O15118 | 557 | Negative | Positive | 0.737 | Positive |
| O15118 | 572 | Negative | Positive | 0.759 | Positive |
| O15118 | 598 | Negative | Positive | 0.847 | Positive |
| O15118 | 916 | Negative | Positive | 0.673 | Positive |
| O15118 | 931 | Negative | Positive | 0.753 | Positive |
| O94923 | 303 | Negative | Positive | 0.665 | Positive |
| O94923 | 393 | Negative | Positive | 0.321 | Negative |
| P00736 | 581 | Negative | Positive | 0.792 | Positive |
| P05106 | 478 | Negative | Positive | 0.937 | Positive |
| P07911 | 275 | Negative | Positive | 0.857 | Positive |
| P07911 | 513 | Negative | Positive | 0.762 | Positive |
| P08887 | 245 | Negative | Positive | 0.871 | Positive |
| P08887 | 350 | Negative | Positive | 0.735 | Positive |
| P09172 | 344 | Negative | Positive | 0.619 | Positive |
| P09172 | 566 | Negative | Positive | 0.793 | Positive |
| P17405 | 505 | Negative | Positive | 0.688 | Positive |
| P38435 | 570 | Negative | Positive | 0.743 | Positive |
| P38435 | 605 | Negative | Positive | 0.900 | Positive |
| P55058 | 117 | Negative | Positive | 0.714 | Positive |
| Q15043 | 87 | Negative | Positive | 0.673 | Positive |
| Q6NSJ0 | 346 | Negative | Positive | 0.797 | Positive |
| Q6NSJ0 | 372 | Negative | Positive | 0.677 | Positive |
| Q6NSJ0 | 398 | Negative | Positive | 0.746 | Positive |
| Q6P179 | 219 | Negative | Positive | 0.754 | Positive |
| Q8IV08 | 236 | Negative | Positive | 0.687 | Positive |
| Q8IV08 | 284 | Negative | Positive | 0.710 | Positive |
| Q8IV08 | 387 | Negative | Positive | 0.848 | Positive |
| Q8N766 | 370 | Negative | Positive | 0.681 | Positive |
| Q8N766 | 818 | Negative | Positive | 0.747 | Positive |
| Q9NV96 | 107 | Negative | Positive | 0.763 | Positive |
| P01266 | 110 | Negative | Positive | 0.730 | Positive |
| P01266 | 1869 | Negative | Positive | 0.854 | Positive |
| P01266 | 2122 | Negative | Positive | 0.691 | Positive |

**Table S4.** Results of MaskNGly predictions of *N*-glycosylation sites for a possible false-positive dataset with revised annotations. These sites were previously annotated as positive but these annotations were removed in the later release. MaskNGly predictions were assigned using a score threshold of 0.5. "Previous" and "Revised" denote annotations based on UniProt releases 2016_08 and 2026_02, respectively.

| UID | Site | Previous | Revised | MaskNGly score | Prediction |
|---|---|---|---|---|---|
| A6NMZ7 | 298 | Positive | Unknown | 0.239 | Negative |
| O75144 | 102 | Positive | Unknown | 0.713 | Positive |
| P10451 | 79 | Positive | Unknown | 0.331 | Negative |
| P10451 | 106 | Positive | Unknown | 0.243 | Negative |
| P19823 | 96 | Positive | Unknown | 0.518 | Positive |
| P27169 | 227 | Positive | Unknown | 0.241 | Negative |
| P47989 | 1074 | Positive | Unknown | 0.320 | Negative |
| Q14517 | 3415 | Positive | Unknown | 0.276 | Negative |
| Q6ZSG1 | 199 | Positive | Unknown | 0.406 | Negative |

### Supplementary Figures

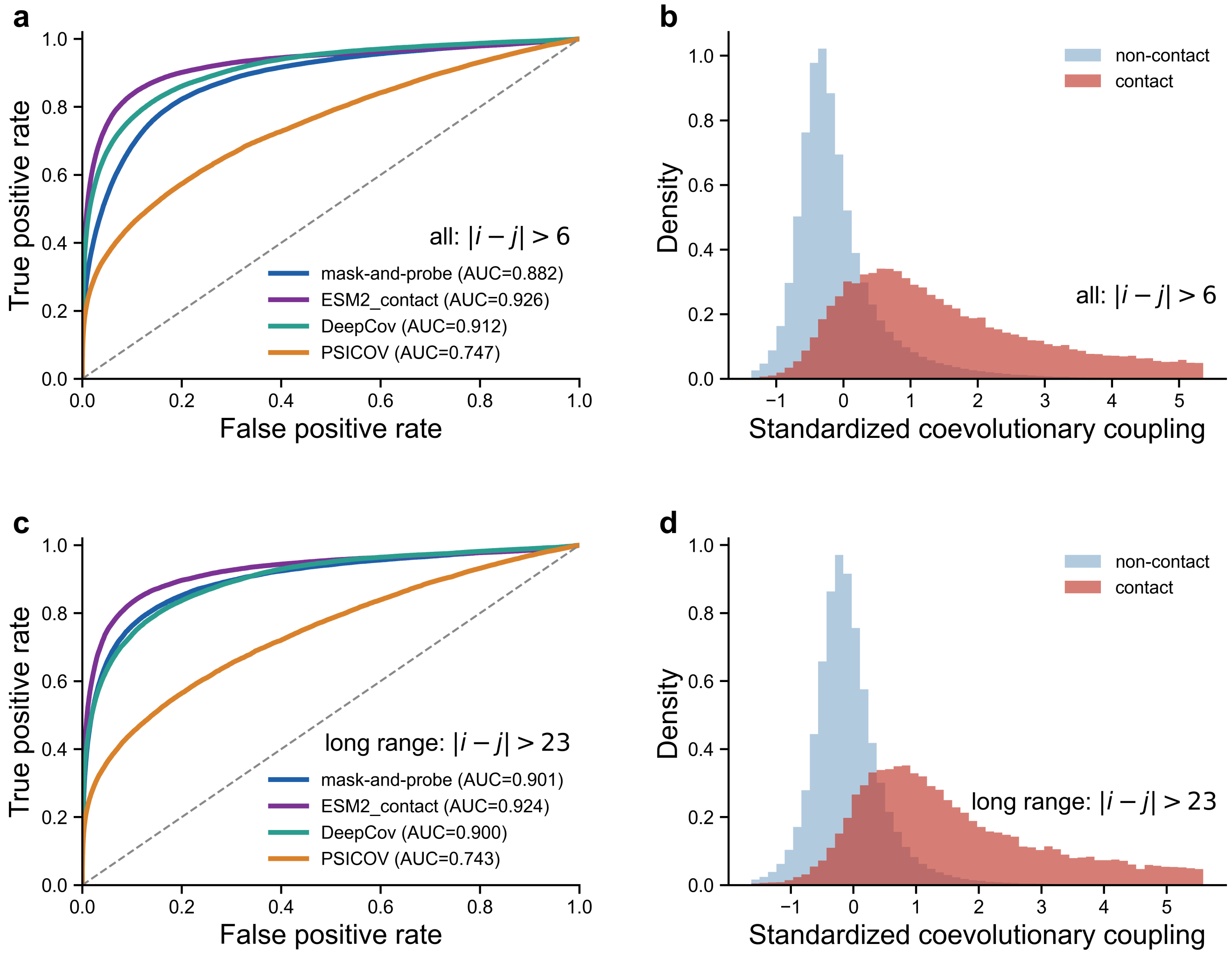


**Figure S1.** Performance comparison among mask-and-probe based coevolutionary analysis (mask-and-probe), DeepCov, PSICOV, and the ESM-2 contact-prediction module. (a,c) Receiver operating characteristic curves (ROC) for the contact predictions using four methods for all residue contacts (a) and for long-range contacts with large sequence separations (c) in the PSICOV150 dataset. The corresponding areas under the receiver operating characteristic curve (AUCs) are indicated in the legends. ROC curves were generated by applying varying thresholds to the coevolutionary-coupling scores when assigning predicted residue–residue contacts. (b,d) Distributions of coevolutionary coupling scores for all residue pairs forming native contacts (b) and for residue pairs forming long-range contacts (d) computed across proteins in the PSICOV150 dataset. For comparison, the corresponding distributions for the residue pairs without forming contacts are also shown (non-contact). The coevolutionary coupling matrices are symmetrized and standardized to zero mean and unit variance for each protein.

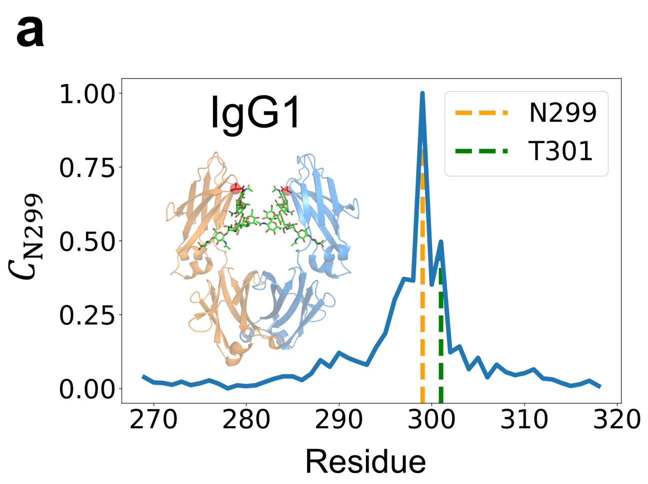


**Figure S3.** The coevolutionary coupling profiles for proteins IgG1 (a) and RecA (b) by masking N299 and N333, respectively.

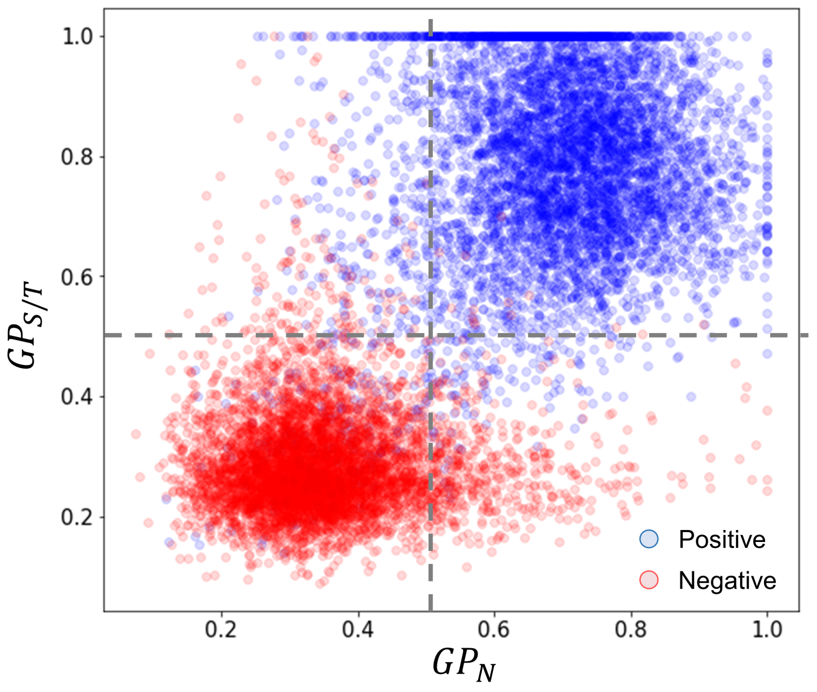


**Figure S4.** Two-dimensional scatter plot of the $GP_N$ and $GP_{S/T}$ scores for the UniProt-derived positive set and the non-glycosylated negative set. Positive samples cluster in the upper-right quadrant ($GP_N > 0.5$, $GP_{S/T} > 0.5$), while negative samples concentrate in the lower-left region ($GP_N < 0.5$, $GP_{S/T} < 0.5$), indicating clear separation between the two groups.

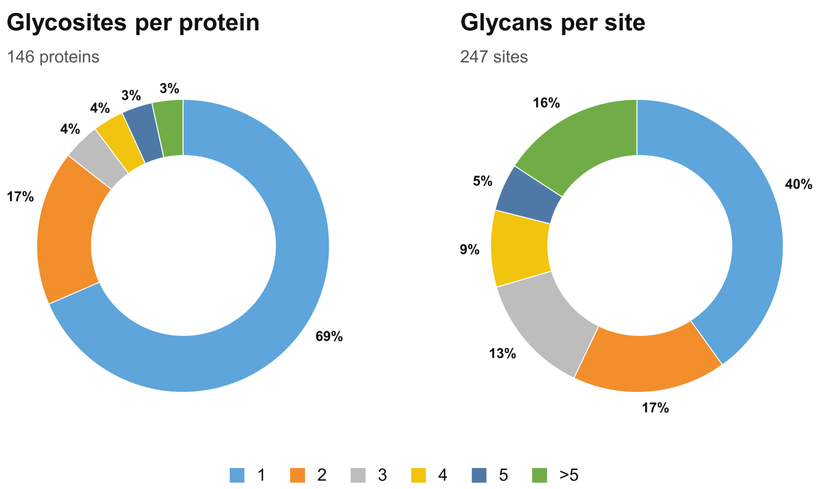


**Figure S5.** Distributions of *N*-glycosylation sites per glycoprotein (left) and glycan composition types per site (right) identified in the glycoproteomic experiment of this study.

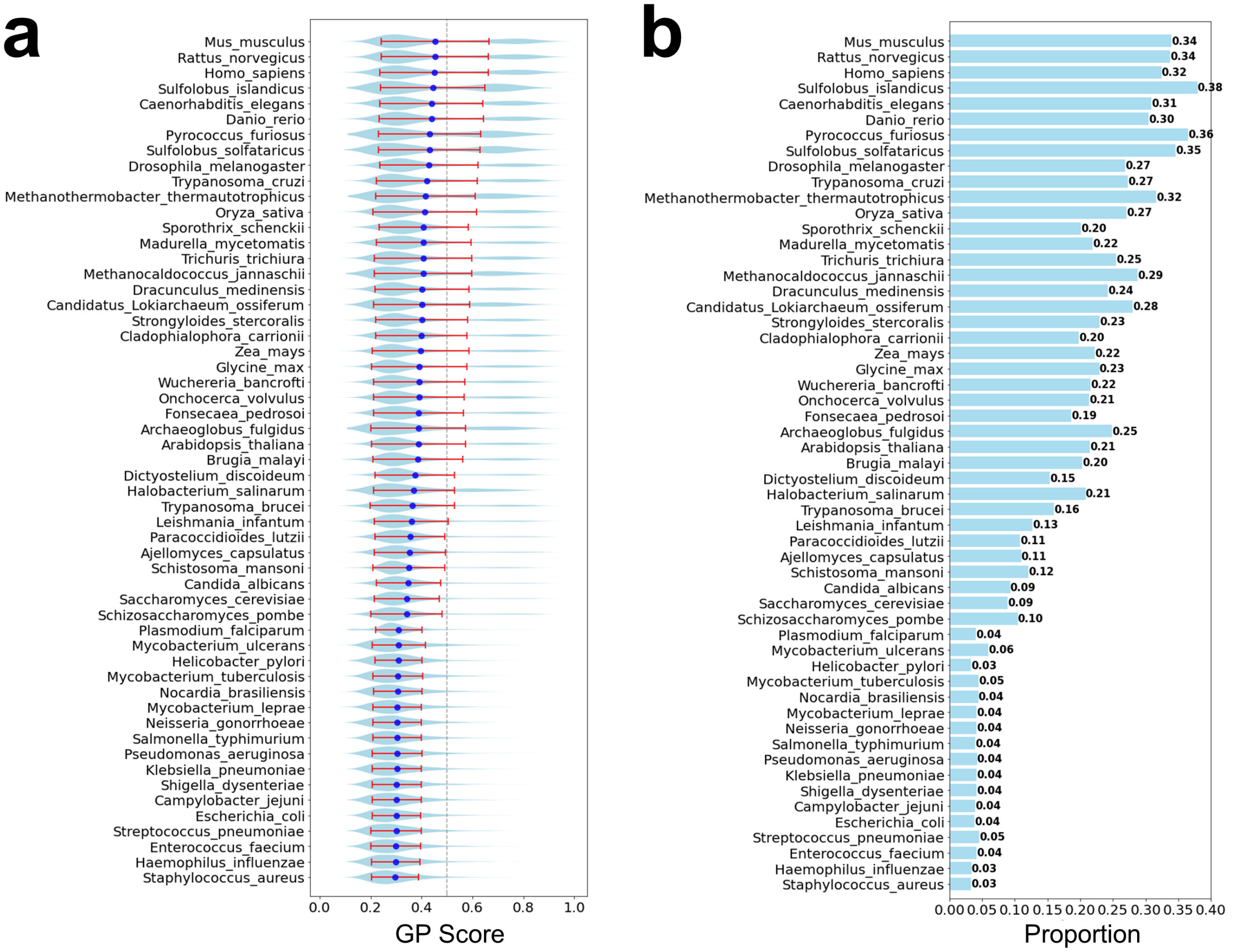


**Figure S8.** Proteome-scale cross-species prediction of *N*-glycosylation sites using sequences from AlphaFold DB and NCBI Datasets [7–9]. (a) Mean prediction scores for different species, with the error bars representing the standard deviation. The distribution of prediction scores is presented with violin diagrams. (b) Proportion of predicted sites with scores above the 0.5 threshold for different species.

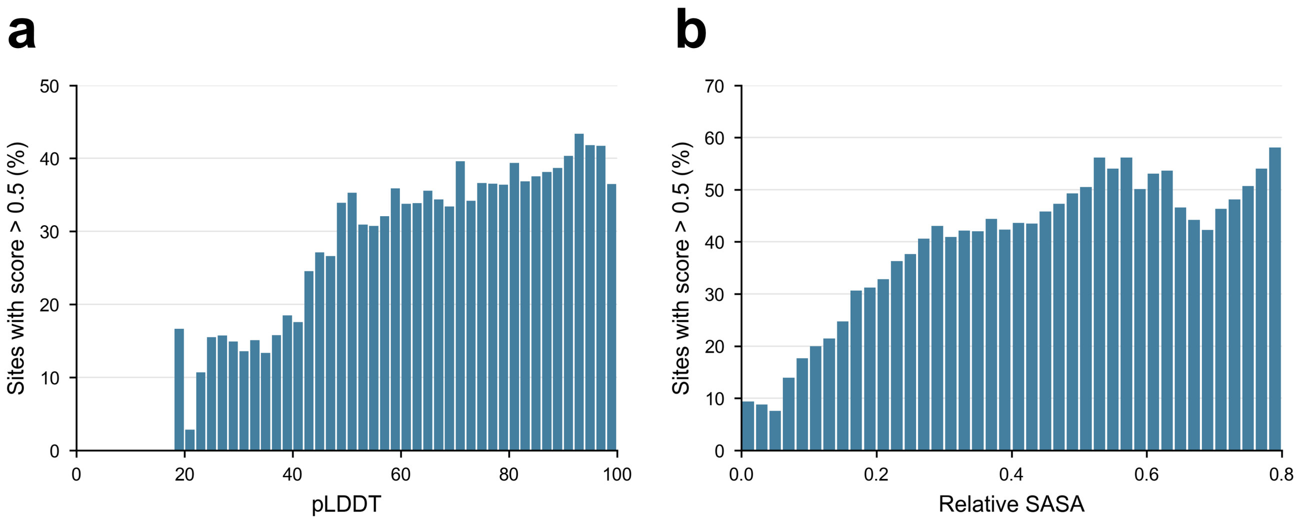


**Figure S9.** Percentages of high-scoring sites across different ranges of pLDDT (a) and relative SASA (b) in human proteins.

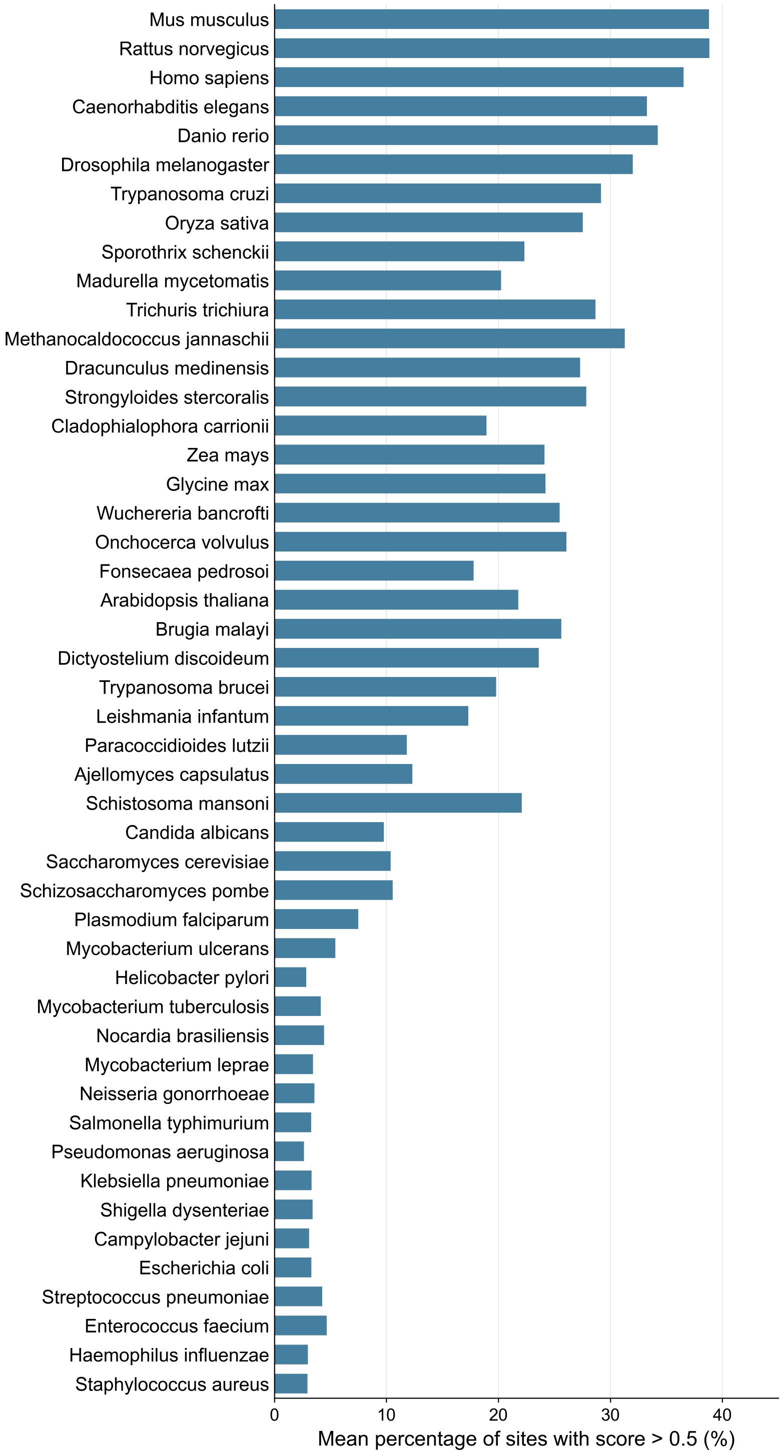


**Figure S10.** Within high-pLDDT regions (pLDDT $>$ 50), the percentage of high-scoring sequons across proteins from different species. Species are ordered as in Fig S8a.

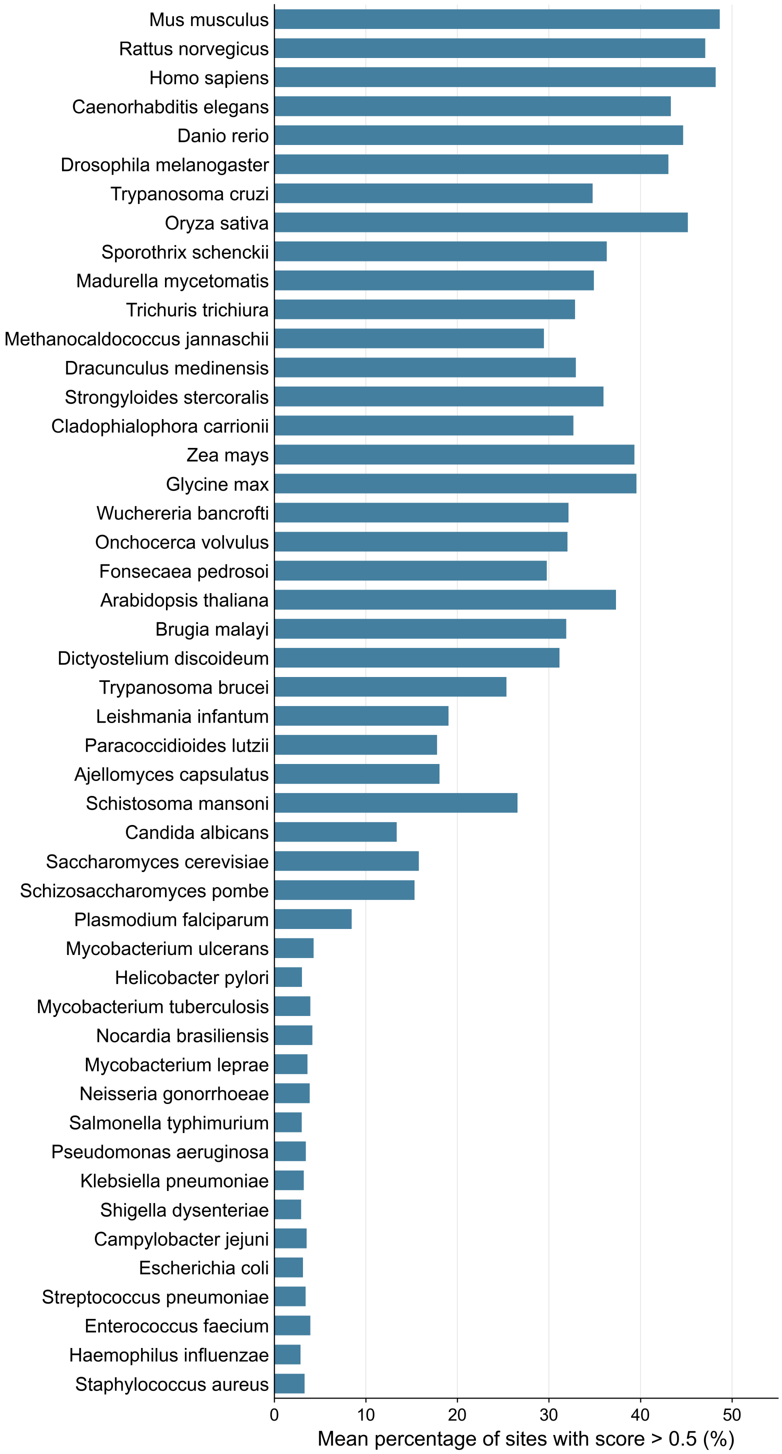


**Figure S11.** Within high-exposure regions (rSASA $> 0.3$), the percentage of high-scoring sequons across proteins from different species. Species are ordered as in Fig S8a.
